# Regulatory elements of pancreas development license the initiation of pancreatic ductal adenocarcinoma

**DOI:** 10.1101/2025.06.06.658264

**Authors:** Marta Ballester, Anita Kurilla, Yifan Dai, H. Carlo Maurer, Elisa Espinet, Kristina Høj, Aida Marisch Delgado, Philip A. Seymour, Saynab Omar, Charlotte Vestrup Rift, Jane Hasselby, Pia Klausen, Peter Vilmann, Albin Sandelin, Luis Arnes

## Abstract

Cellular plasticity and transitional cellular states are crucial for tissue regeneration across multiple organs. In the pancreas, oncogenic Kras hijacks this program, acting on tissue-specific enhancers to prevent the resolution of acinar-to-ductal metaplasia (ADM) and lock regeneration into a pro-inflammatory state that progresses to cancer. Enhancer transcription, an early event during cellular state transitions, can generate stable enhancer-associated long noncoding RNAs (lncRNAs) positioned near key transcription factors and chromatin contact boundaries, often enriched for disease-associated variants. While enhancer-associated lncRNAs have been implicated in transcriptional regulation and genome organization, their role in pancreas regeneration and cancer initiation has remained unexplored.

In this study, we investigated the expression of epithelial long noncoding RNAs (lncRNAs) and their target genes in PDAC precursor lesion formation. We focus on lncRNAs transcribed from enhancer elements near cell identity transcription factors. We demonstrate that LINC00673, expressed from a Sox9-associated super-enhancer during pancreatic development, is reactivated in PDAC. Conditional deletion of LINC00673 in the murine pancreatic epithelium accelerates resolution of ADM and significantly impairs PDAC initiation. Notably, LINC00673 harbors a variant associated with risk of developing PDAC. Our study identifies a critical function of LINC00673 in regulating both cell-autonomous and non-cell-autonomous processes during pancreas regeneration and Kras-driven cancer initiation. Furthermore, we highlight a previously unrecognized role of transcribed super-enhancers in facilitating long-range gene regulation during pancreatic cancer initiation. These findings reveal a novel regulatory layer linking developmental enhancer activity, cellular plasticity and pancreatic disease progression.

**Teaser:** Long noncoding RNAs from developmental enhancers play a role in long-range gene regulation with crucial biological impacts in development and cancer.

**Grant support:** This project has been supported by the Danish Cancer Society (R302-A17481). LA is supported by core funding of the Biotech Research and Innovation Center, the Danish Cancer Society (R302-A17481, R322-A17.350), The Novo Nordisk Foundation (NNF21OC0070884) and The Innovation Fund (Eurostars 2807). The Novo Nordisk Foundation Center for Stem Cell Biology was supported by Novo Nordisk Foundation grants NNF17CC0027852.

## Introduction

Tissue injury increases the risk of developing pancreatic ductal adenocarcinoma (PDAC) through mechanisms that remain poorly understood. Upon damage, acinar cells adopt a ductal phenotype through the silencing of markers of differentiation and re-initiation of pancreas developmental programs ^1–3^. This mechanism, termed acinar-to-ductal metaplasia (ADM), alleviates tissue damage, induces robust acinar proliferation, and is necessary for regeneration ^4^. However, in the event of a mutation in the *Kirsten Rat Sarcoma* (Kras) oncogene, ADM lesions are susceptible to malignant transformation through formation of pancreatic intraductal neoplasia (PanIN) ^5,6^. In comparison, differentiated acinar cells are protected from neoplastic transformation but their regenerative capacity is limited to tissue turnover in homeostasis ^7^. These findings emphasize the importance of understanding the regulation of transient cellular states in the adaptation to tissue injury, the resolution of the regenerative program, and the interaction between developmental programs and genetic alterations in the early stages of PDAC.

Transient cellular states, common to regenerative programs in multiple tissues, depend on the reactivation of developmental enhancers active in tissue related developmental contexts^8^. Oncogenes, particularly *Kras*, exploit this inherent plasticity by hijacking tissue-specific enhancers, thereby preventing the resolution of ADM and locking pancreas regeneration in a persistent pro-inflammatory phase that predisposes tissue to cancer^9^. Transcription is widespread at mammalian enhancers, and represents the first wave of genome activity during differentiation and cellular state transitions (reviewed in ^10^). In certain contexts, enhancer transcription generates stable and evolutionarily conserved enhancer-associated long noncoding RNAs (lncRNAs) ^11,12^. These lncRNAs are often located near transcription factors and at boundaries of three-dimensional chromatin contacts, and they are enriched in disease-associated genetic variants ^11–14^. Such observations suggest critical regulatory roles for enhancer-associated lncRNAs in transcriptional control, genome organization, and cell-fate decisions across diverse biological processes. However, the contribution of developmental enhancers to pancreatic regeneration and cancer initiation remains unexplored, and therapeutic strategies targeting regenerative programs remain underdeveloped.

To explore this further, we focused on one particular enhancer-associated lncRNA, LINC00673, which is transcribed from a developmental super-enhancer in the SOX9 locus and reactivated during pancreatic injury and neoplastic transformation. Our study sheds light on the mechanisms by which enhancer-associated lncRNA regulate cell fate decisions in pancreas regeneration and cancer initiation. We reveal a novel layer of gene regulation essential for Kras-mediated cancer initiation, involving both cell-autonomous and non-cell-autonomous processes in early preneoplastic lesions. These insights advance our understanding of the complex interplay between transient epithelial cellular states, the tissue microenvironment, and pancreatic regeneration and cancer.

## Results

### Gene regulatory network analysis identifies lncRNA regulators associated with cancer initiation

In this study, our objective was to identify transcribed regulatory elements of pancreas development reactivated in transitional cellular states in pancreas regeneration and investigate their role in the onset of PDAC. To achieve this goal, we employed gene expression profiles from benign precursor lesions and primary PDAC human specimens. Our cohort consisted of 242 transcriptional profiles derived from laser capture microdissected epithelium (Figure 1A, data from ^15^). We reverse-engineered a gene regulatory network using the ARACNe (Algorithm for the Reconstruction of Accurate Cellular Networks)^16^. To focus on lncRNAs expressed from potential enhancers of transcription factors, we considered lncRNAs that (i) exhibited sufficient expression levels in epithelial samples and (ii) were located within 200 kb of a transcription factor gene. This analysis revealed a network composed of 1,127 lncRNAs and 16,520 target genes, comprising 76,794 interactions (Supplementary Table 1). Importantly, during network engineering, we allowed transcription factors, alongside lncRNAs, to serve as regulatory nodes. This ensured that only genes without stronger mutual information with a transcription factor were retained as lncRNA targets. Notably, the analysis identified lncRNAs located near transcription factors critical for pancreas development and PDAC progression, including GATA6, HNF1A, SOX9, and FOXA2 ^17–20^. Gene ontology analysis of the target genes revealed significantly enriched categories such as "DEVELOPMENTAL PROCESSES," "CELL MIGRATION," "HYPOXIA," and "PANCREATIC CANCER SUBTYPES" (Supplementary Table 2), supporting a potential role for lncRNAs in the malignant transformation of epithelial cells during PDAC development.

**Figure 1:**
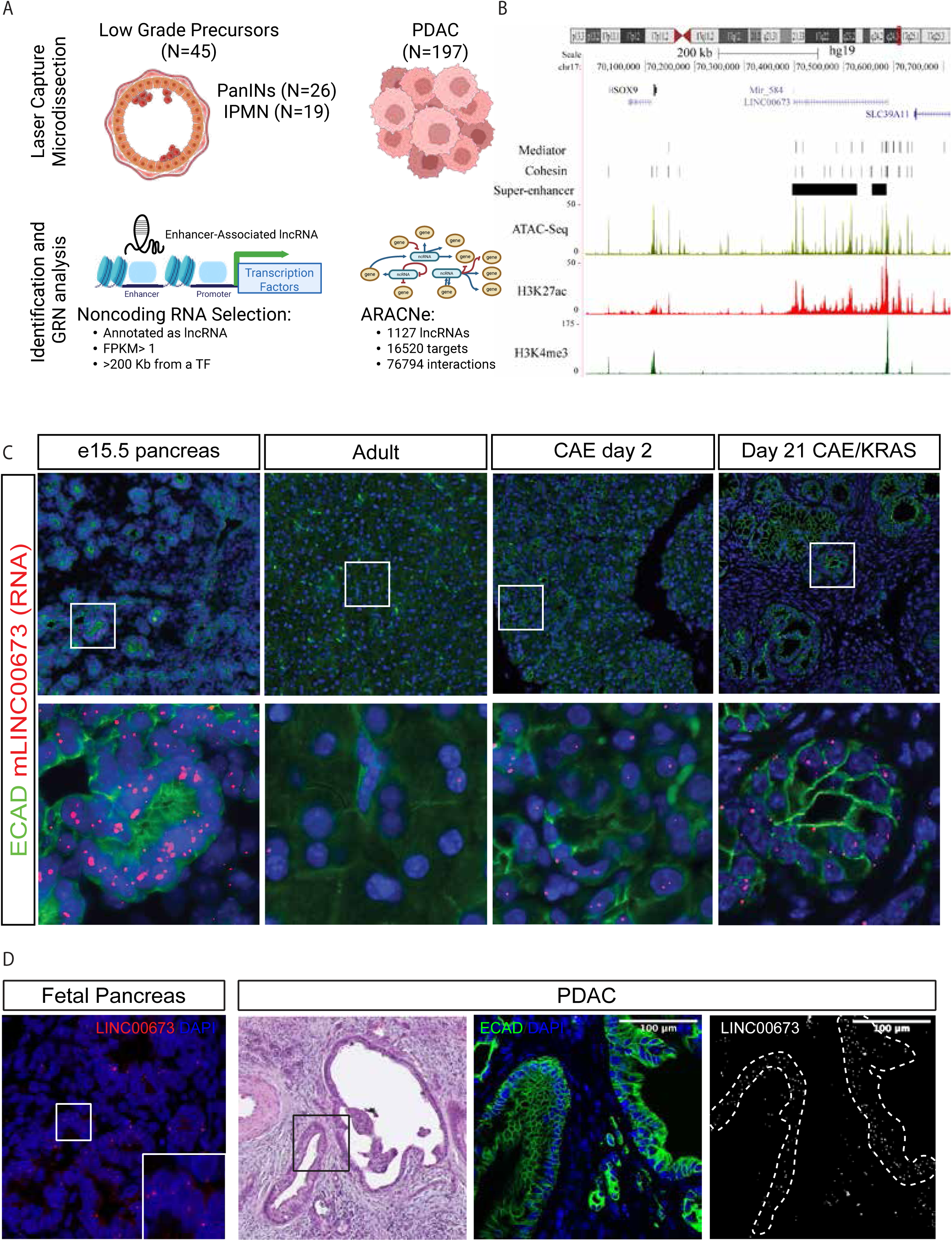
Gene regulatory network of cancer initiation identifies the lncRNA LINC00673 transcribed from a super-enhancer. A) To identify lncRNAs involved in PDAC progression from precursor lesions, we used the ARACNe algorithm on RNA sequencing data of laser micro dissected human samples of precursor lesions (n=45) and established PDAC (n=197). Data from Maurer et al., bioRxiv 2024. B) UCSC Genome browser snapshot of the Sox9-LINC00673 locus. The LINC00673 locus is enriched in binding sites of cohesins (Smca1 and Med1) in PANC-1 cells and is annotated as a super-enhancer in islets of Langerhans ^24^. C) mLINC00673 is expressed in development (e15.5 pancreas) and reactivated upon tissue injury (CAE day 2) and in preneoplastic lesions of PDAC (Day 21 CAE/KRAS) seen by single molecule in situ hybridization (RNAscope). Representative images of at least three animals. D) Left: we performed RNAscope of human pancreatic embryonic tissue 7 weeks from conception. Right; H&E and ISH coupled with IF (Krt7, green) of LINC00673 of human tumor sample. LINC00673 expression is restricted to the tumor epithelium (white dotted line). Representative images of at least three tissue samples.

Among the identified lncRNAs, LINC00673 emerged as a particularly interesting candidate. It is located within a gene desert downstream of SOX9, an essential transcription factor in pancreas development and cancer initiation (Figure 1B). Furthermore, the LINC00673 locus harbors a single nucleotide polymorphism (SNP) associated with increased PDAC risk, as identified in genome-wide association studies ^21^. While most regulatory elements of SOX9 have been characterized upstream of the gene, very little is known about the regulatory landscape downstream, highlighting a potential novel regulatory mechanism involving LINC00673.

LINC00673 is conserved across tetrapods ^22^. In-depth analysis of epigenetic marks in the LINC00673 locus revealed chromatin modifications associated with a transcribed super-enhancer, along with binding sites for key factors such as CTCF, cohesins, BRD4, the mediator complex and transcription factors important in pancreas development and malignant transformation (Figure 1B, Supplementary Figure 1A) ^23,24^. The mouse ortholog, 2610035D17Rik (hereafter referred to as *mLINC00673*), has an 83.17% identity with human LINC00673 and is conserved in synteny in mouse and human. These findings suggest that the LINC00673 locus serves as a conserved regulatory hub, integrating structural and transcriptional control functions essential for genome regulation.

### LINC00673 is dynamically expressed during pancreas development and reactivated in pancreas regeneration and cancer

To investigate the LINC00673 pattern of expression in pancreas development, regeneration, and cancer, we used an experimental model of tissue injury and cancer initiation. We induced experimental pancreatitis in the *Pdx1-Cre; Kras*^LSL-G12D*/+*^*; Rosa26*^LSL-YFP/LSL-YFP^ (KCY) mouse model ^25,26^ through the repeated administration of a supraphysiological level of the cholecystokinin analogue, caerulein (CAE). While the activation of KRASG12D is weakly oncogenic in homeostasis, it synergizes with signals activated upon tissue injury and inflammation to generate PanINs^26^. To determine the expression of *mLINC00673* and its cellular localization in mouse, we performed single-molecule RNA *in situ* hybridization (RNAscope). We observed *mLINC00673* expression in the epithelium of the developing pancreas at E15.5 during the peak of the secondary transition when differentiation into endocrine, ductal and acinar cells is maximal during pancreogenesis (Figure 1C, Supplementary Figure 2A). While *mLINC00673* expression was hardly detectable in the postnatal pancreas under homeostatic conditions, it exhibited a pronounced upregulation in the pancreatic epithelium following the induction of tissue injury, particularly with the concomitant expression of mutant Kras (Figure 1C). *mLINC00673* expression persisted across the majority of epithelial cells in PanINs (Figure 1C). Remarkably, subcellular localization of *mLINC00673* expression is mostly nuclear and the transcripts in the nucleus appeared to be predominantly associated with two discrete structures. This observation suggests that the molecular function of *mLINC00673* may be restricted to the nuclear compartment in mice. Transcriptome analysis confirmed this dynamic expression pattern during pancreas development and cancer initiation (Supplementary Figure 2B). Furthermore, analysis of chromatin accessibility and expression in sorted acinar cells showed that whereas the locus becomes accessible upon injury, the expression is significantly upregulated only in the presence of mutant Kras activation (Supplementary Figure 2C-D)^27^.

Next, we investigated whether the pattern of expression of LINC00673 in pancreas development, regeneration and cancer was conserved in human samples. We observed LINC00673 expression in the epithelium of developing pancreas and in tumor cells from primary PDAC (Figure 1D). To acquire more comprehensive insight into the pattern of expression in development and cancer, we took advantage of human pluripotent stem cells (hPSC), organoids derived from primary PDAC tumors and public dataset of chronic pancreatitis (Supplementary Figure 3). Using a hPSC pancreatic differentiation protocol that recapitulates elements of pancreas development ^28^, we determined that LINC00673 expression peaks upon the emergence of pancreatic progenitors (Supplementary Figure 3A). To assess whether LINC00673 expression is activated upon tissue injury, we exploited single-nucleus RNA sequencing from human chronic pancreatitis samples and identified LINC00673 expression in a subpopulation of MUC5ACB+/KLF5+ cells, reminiscent of the transitional cellular state observed in ADM and PanINs (Supplementary Figure 3B)^30^. Moreover, we determined that LINC00673 expression is associated with the classical molecular subtype of PDAC using gene expression analysis in sorted epithelial cells from resected tumors (Supplementary Figure 3C) ^31^. Accordingly, expression of LINC00673 assessed by ISH was found in GATA6+ tumor-derived organoids (Supplementary Figure 3D) associated with the classical molecular subtype. Taken together, these results demonstrate that *mLINC00673* expression is dynamically regulated during development, reactivated in a transitional cellular state of acinar dedifferentiation that emerges following tissue injury, and is maintained in precancerous lesions of PDAC and early stages of tumor progression in both mice and humans.

### Genetic inactivation of mLINC00673 protects against KrasG12D-induced PanIN formation in vivo

Next, we sought to investigate the function of LINC00673 in pancreas regeneration and cancer using mouse genetics. We generated mice in which loxP sites flank exon 3 of *mLINC00673 (mLINC00673-E3)* using BAC recombineering (Supplementary Figure 4A; Materials and Methods). The resulting *mLINC00673-E3-flox* mice were crossed with *Pdx1-Cre* to delete *mLINC00673-E3* specifically in the pancreatic epithelium and all cellular progeny during the early stages of pancreas development. We generated *Pdx1-Cre; mLINC00673-E3^fl/fl^* (CL, mLINC00673-E3 cKO) and *Pdx1-Cre; mLINC00673^+/+^* (C) mice (Supplementary Figure 4B). This strategy allowed us to target the longest exon of LINC00673 that in humans harbors a SNP associated with PDAC risk (rs11655237) ^32^ while preserving the transcription start site containing epigenetic marks of active transcription, enhancer and transcription factor (Supplementary Figure 1B).

Mice carrying either one or two copies of the loxP allele were born in Mendelian ratios and did not exhibit any morphological defects in the pancreas (*data not shown*). Conditional deletion of mLINC00673-E3 in the developing pancreas did not result in morphological abnormalities (Supplementary Figure 5A). However, male mice exhibited mild glucose intolerance by 20 weeks of age (Supplementary Figure 5B). To confirm the deletion of *mLINC00673* we performed quantitative PCR on E15.5 embryonic pancreata which showed efficient deletion in the developing pancreas in CL mice compared to C controls (Supplementary Figure 5C). Notably, the expression of exon 1 was not affected by exon 3 deletion, suggesting that the *mLINC00673* promoter and transcription were not affected by the genetic manipulation (Supplementary Figure 5C).

Next, we aimed to investigate the function of mLINC00673 in the adaptation of acinar cells to injury and cancer initiation. We assessed the role of *mLINC00673* in the formation of PanIN lesions following tissue injury and KRAS mutation. We generated mice that combined KrasG12D activating mutations with the deletion of *mLINC00673* in the pancreatic epithelium. Specifically, we generated two groups of mice: *Pdx1*-*Cre; Kras^LSL-G12D/+^; mLINC00673^+/+^* (KC), *Pdx1-Cre; Kras^LSL-G12D/+^; mLINC00673^fl/+^* (KCL^HET^), and *Pdx1-Cre; Kras^LSL-G12D/+^; mLINC00673^fl/fl^* (KCL^HOM^). We induced tissue injury in 8-week-old mice and collected pancreata after 21 days for further analysis (Figure 2A). We validated *mLINC00673* recombination by RNAscope, revealing a complete absence of *mLINC00673* transcript in many epithelial cells in KCL^HOM^ mice compared to expected upregulation in the KC mice, indicating efficient recombination in the pancreatic epithelium (Supplementary Figure 6). Histological analysis revealed morphological evidence of Krt19+ ductal-like lesions, disturbed lobular architecture, extensive fibrosis, large irregular ductal structures lined by cells with large and hyperchromatic nuclei in KC, but these features were greatly reduced in KCL mice (Figure 2B). Morphometric analysis of lesions showed an almost complete depletion of PanINs in pancreata from KCL^HOM^ compared to KC mice (Figure 2B). The reduction of PanINs was also associated with a significant decrease in the desmoplastic reaction, evident by a lower number of both cells expressing Vimentin (Vim) (Figure 2C) and F4/80+ macrophages (Figure 2E). The reduced burden of PanIN lesions was associated with an increased survival of KCL^HOM^ mice compared to KC controls (Figure 2D). These results suggest that mLINC00673 is necessary for induction of PDAC precursors and the remodelling of the associated premalignant tissue microenvironment.

**Figure 2:**
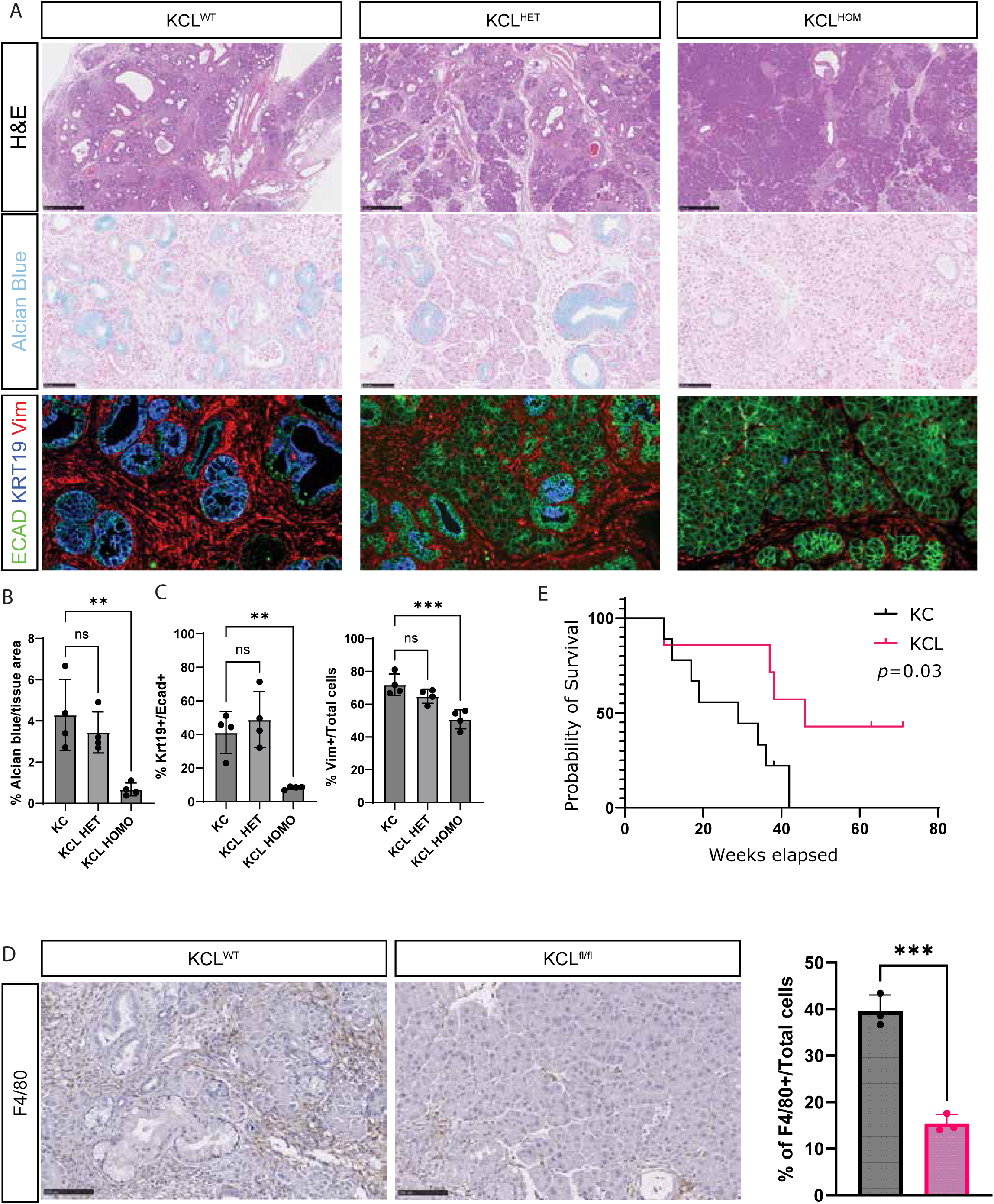
mLINC00673 potentiates the initiation of pancreatic cancer. A) Histological analysis of pancreata 21 days after caerulein-induced pancreatitis in Pdx1-Cre; Kras^LSL-G12D/+^; mLINC00673^+/+^ (KC), Pdx1-Cre; Kras^LSL-G12D/+^; mLINC00673^f/+^ (KCL^HET^), and Pdx1-Cre; Kras^LSL-G12D/+^; mLINC00673^fl/fl^ (KCL^HOM^) mice. (B) Alcian blue staining revealed a loss of mucin-producing cells in KCL^HOM^ compared to KC and KCL^HET^. (C) Immunofluorescence showed reduced Krt19^+ lesions and fewer Vim+ stromal cells in KCL^HOM^ mice. Quantification includes alcian blue+ cells, Krt19+/Ecad+ epithelial cells, and Vim+ stromal cells (n = 4 mice per group). (D) F4/80 immunohistochemistry revealed abundant macrophages in KC PanIN lesions, which were reduced in mLINC00673-deficient mice showing restored acinar morphology. Right, quantification of F4/80^+ cells. (E) Kaplan-Meier survival analysis of KC (n = 9) and KCL (n = 7) mice (*p = 0.03, log-rank). *p < 0.05, **p < 0.01, ***p < 0.001 versus KC. Data represent mean ± SD.

Given that non-pancreatic tissues of endodermal origin have common regulatory elements in development, we asked whether these observations could be extrapolated to other endoderm-derived tumors. We took advantage of publicly available forward genetic screenings in genetically engineered mouse models of cancer ^33^. Supporting our observations in cancer initiation, *mLINC00673* is significantly enriched in Sleeping Beauty transposon insertional mutagenesis in genetic mouse models of primary PDAC (Supplementary Figure 7) ^34^. Strikingly, *mLINC00673* is also represented in endoderm-derived tumors driven by mutations in Kras, Smad4, WNT signaling and Trp53 (e.g., stomach and intestine) and not in tumors arising from tissues derived from the other germline layers (e.g., leukemia and skin cancer). Our data demonstrate that *mLINC00673* is necessary for the formation of PanINs and suggest that it regulates the progression of endoderm-derived tumors, including those of the gastrointestinal epithelium.

### mLINC00673 is required for stromal remodeling but is dispensable for acinar plasticity in KRAS-driven pancreatic lesions

PanINs originate from the interaction of transcriptional programs of pancreas regeneration with oncogenic KRAS, which locks the regenerative program of acinar-derived transitional cellular states into a profibroinflammatory state ^2,27,35,36^. To investigate the function of *mLINC00673* in the early events leading to tissue remodeling and neoplastic transformation, we subjected KC/KCL^HOM^ and C/CL mice to CAE-induced pancreatitis and harvested tissues two days after treatment.

Histopathological examination of KC and KCL^HOM^ pancreata revealed comparable morphological features of ADM, indicating that mLINC00673 is not required for acinar plasticity or cellular dedifferentiation. Despite this, we observed a marked reduction in stromal cell infiltration in KCL^HOM^ mice compared to KC controls (Figure 3A). Specifically, the number of αSMA+ (ACTA2+) myofibroblasts (Figure 3B) and F4/80+ macrophages (Figure 3C) was significantly decreased in the absence of mLINC00673, an effect that was evident only in the context of oncogenic KRAS activation. In contrast, both C and CL mice exhibited full histological recovery by day seven post-injury (Supplementary Figure 8A). Overall, our data suggest that mLINC00673 is necessary for tissue microenvironment remodeling in the context of mutant KRAS, and that this function appears to be independent of acinar plasticity.

**Figure 3:**
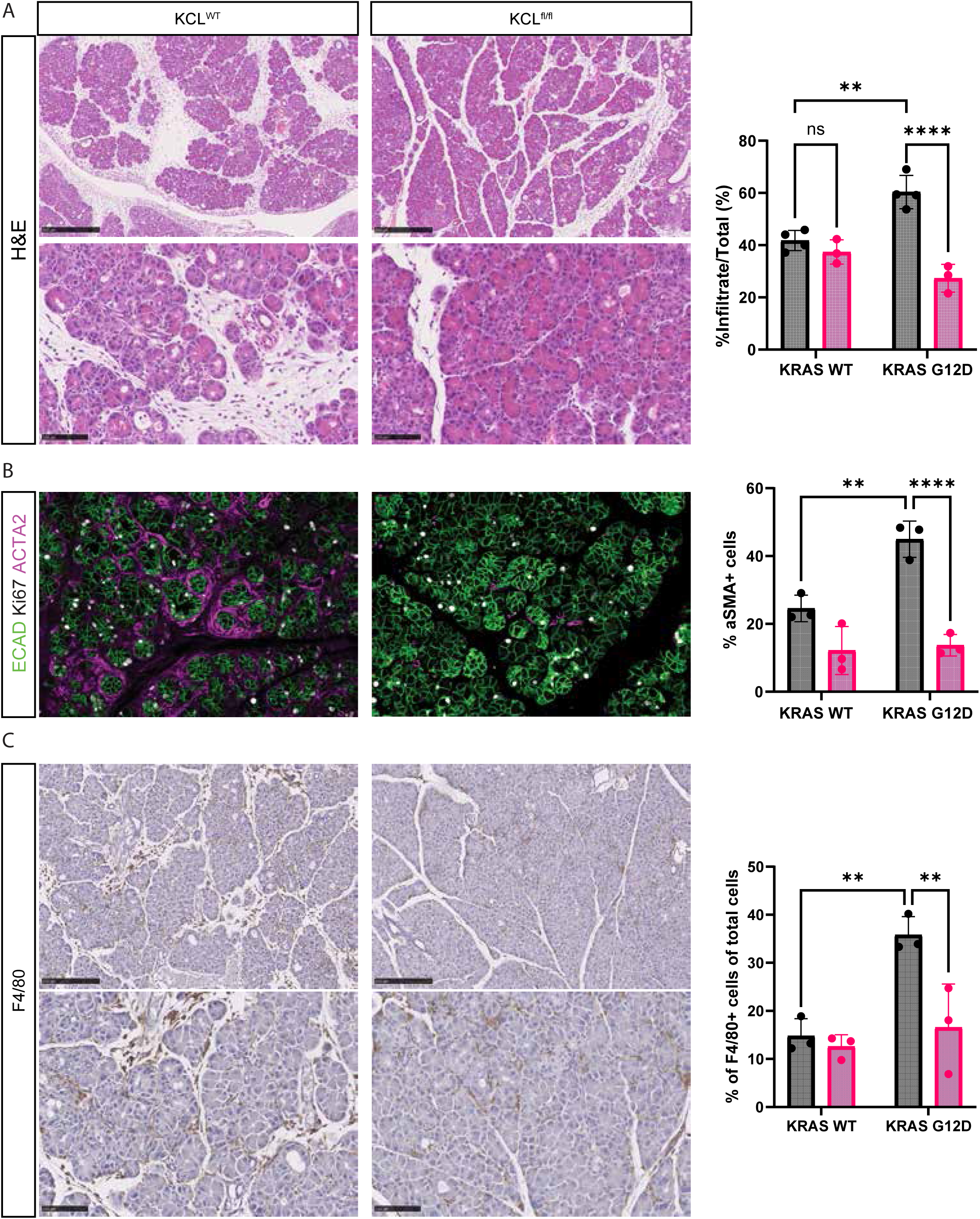
mLINC00673 regulates tissue homeostasis through non-cell autonomous mechanisms. (A–C) Histopathological and immunofluorescence analysis of pancreatic tissue 2 days after caerulein-induced injury in KC and KCL littermates. (A) Evidence of acinar-to-ductal metaplasia (ADM) was observed in both genotypes. Right, quantification of stromal cells as a percentage of total cells (n ≥ 3). (B) KCL mice showed reduced stromal activation and fewer ACTA2+ myofibroblasts compared to KC mice. Right, quantification of ACTA2+ cells (n ≥ 3). (C) F4/80 immunohistochemistry revealed decreased macrophage infiltration in KCL mice. Right, quantification of F4/80+ cells per total cells. Quantification includes CL and C samples (KRAS WT) for reference. *p < 0.05, **p < 0.01, ***p < 0.001 by two-way ANOVA with multiple comparisons. Data represent mean ± SD.

### mLINC00673 is necessary for PanIN formation independently of Sox9 expression

Given the established role of Sox9 in acinar cell dedifferentiation and PanIN development ^17^ and considering that enhancer-associated lncRNAs frequently regulate the transcription of neighboring genes, we assessed Sox9 expression in pancreatic tissues. Despite efficient and near-complete deletion of mLINC00673, nuclear Sox9 expression was readily detectable in epithelial cells of both control C and CL pancreata during regeneration and PanIN formation (Supplementary Figure 9). These findings indicate that mLINC00673 does not regulate Sox9 expression and suggest that its role in PanIN formation is independent of this key transcription factor.

### Loss of mLINC00673 disrupts the immunoinflammatory response in acinar cells

To further elucidate how mLINC00673 influences acinar cell plasticity and oncogenic transformation, we employed a lineage-specific, inducible recombination strategy. Because *Pdx1-Cre* drives recombination throughout the pancreatic epithelium during embryogenesis, we crossed *mLINC00673-E3^fl/fl^* animals with *Kras*^LSL-G12D*/+*^*; Rosa26*^LSL-YFP/LSL-YFP^ mice carrying acinar-specific *Ptf1a*^CreER^ to generate control *Ptf1a^CreER^; Kras*^LSL-G12D/+^*; Rosa26*^LSL-YFP/LSL-YFP^ (KPtC) and *Ptf1a^CreER^; Kras*^LSL-G12D/+^*; Rosa26*^LSL-YFP/LSL-YFP^*; mLINC00673-E3^fl/fl^* (KPtCL) mice. Tamoxifen was administered postnatally to induce recombination in adult, differentiated acinar cells, followed by caerulein to trigger acute pancreatic injury. Tissues were collected at 2 and 21 days post-injury (Supplementary Figure 10A).

Consistent with our observations using the *Pdx1-Cre* driver, *Ptf1a^CreER^*-mediated deletion of mLINC00673 exon 3 delayed the formation of acinar-derived PanIN lesions but did not impair the ability of acinar cells to undergo ADM. Notably, we observed a reduction in inflammation at day 2 post-injury in KPtCL compared with control mice. Collectively, these results indicate that mLINC00673 is not essential for acinar cell dedifferentiation or ADM, but it plays a critical role in promoting the fibro-inflammatory response that accompanies early neoplastic transformation in the pancreas.

### mLINC00673 regulates the expression of genes within a syntenic region in chromosome 11 and 17 in mouse and human respectively

To gain molecular insight into the function of mLINC00673, we examined its subcellular localization. RNAscope revealed that transcripts originating from exon 3 are predominantly retained in the nucleus of both mouse and human cells (Figure 4A). The RNA localized to two discrete nuclear foci within pancreatic progenitors and acinar cells undergoing ADM, suggesting a potential nuclear regulatory role.

**Figure 4:**
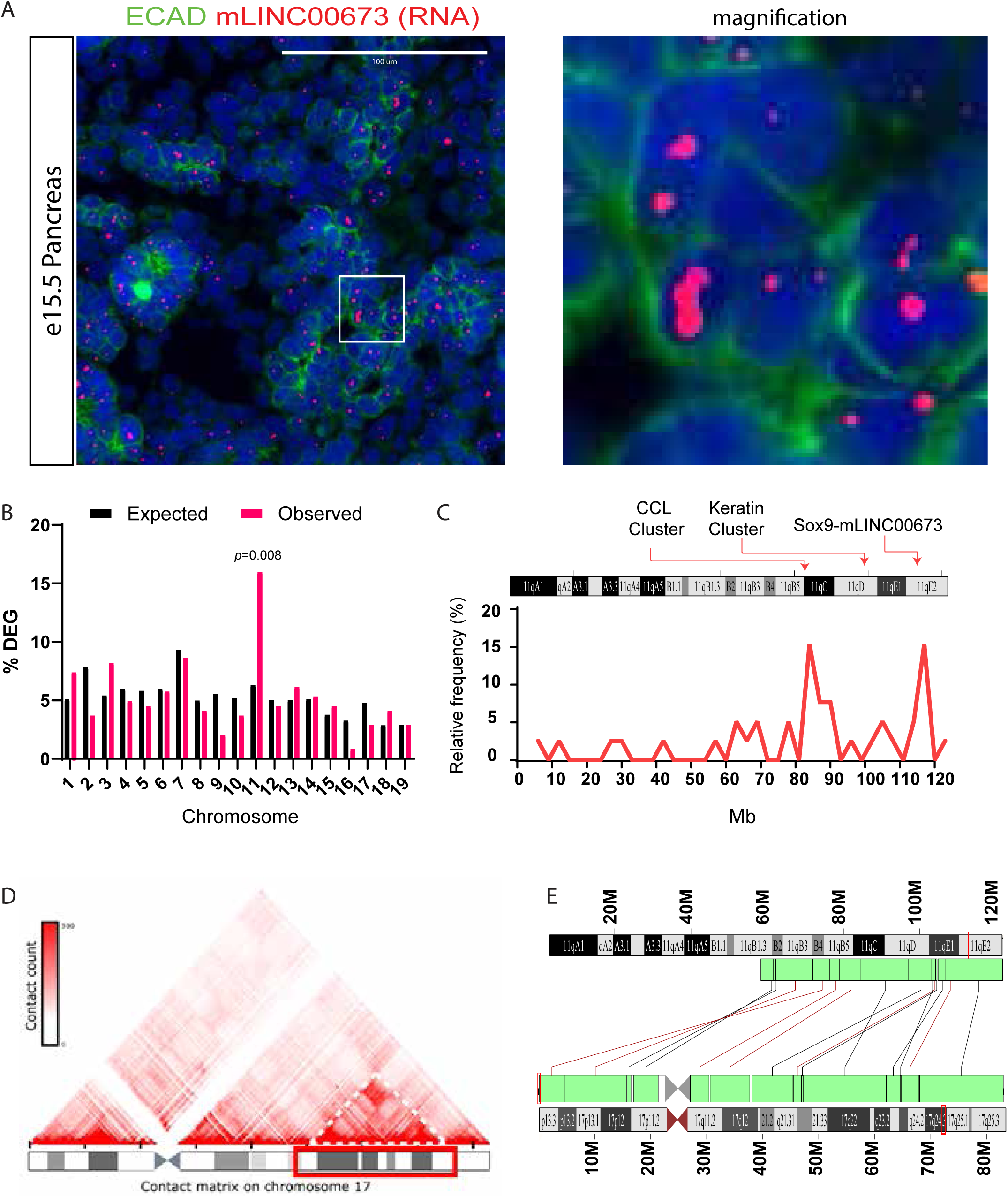
mLINC00673 regulates gene expression in a spatially restricted domain on mouse chromosome 11. (A) RNAscope of E15.5 mouse pancreas shows mLINC00673 RNA (magenta) localized as two nuclear puncta per cell. Representative images from n = 3 mice. (B) RNA-seq and differential expression analysis comparing E15.5 pancreata from mLINC00673^fl/fl^ and Pdx1-Cre; mLINC00673^fl/fl^ embryos revealed chromosome-wide dysregulation. Shown is the proportion of dysregulated genes per chromosome, normalized to total gene count. Loss of mLINC00673 led to significant dysregulation of genes on chromosome 11 (n=3). (C) Density plot of dysregulated genes on chromosome 11 highlights their enrichment in the distal region, syntenic to human chromosome 17. Locations of the C-C motif chemokine ligand cluster, keratin gene cluster, and the Sox9–mLINC00673 locus are indicated. (D) The location of LINC00673 target genes (red box) coincides with a region of frequent chromatin interactions (dotted lines) in Panc1 Hi-C data. (E) Schematic comparison of mouse chromosome 11 and human chromosome 17, showing conserved synteny in the distal region (green blocks, ENSEMBL).

Transcriptomic profiling of the embryonic pancreas at E15.5 identified 240 differentially expressed genes (DEGs, p < 0.05) between CL and control littermates (Supplementary Table 5). To exclude potential confounding effects from loxP site insertion, mLINC00673^fl/fl^ mice were used as controls. Strikingly, many DEGs were clustered within a discrete 20 Mb region in the distal portion of chromosome 11, proximal to the mLINC00673 locus (Figure 4B), suggesting a non-random, spatially restricted regulatory influence. These findings mirror previous observations in which deletion of a super-enhancer near Nkx2-2 similarly affected gene expression within a confined chromosomal domain ^37^.

To assess the functional relevance of genes within this conserved region, we examined their potential involvement in PDAC initiation. Several gene clusters implicated in tumor progression were found within the region, including: CCL chemokines (CCL2, CCL8, CCL9), which modulate the immune microenvironment ^38,39^; RNF43, a negative regulator of Wnt signaling frequently mutated in PDAC precursors ^40^; and a keratin cluster (KRT17, KRT19, KRT7), re-expressed during ADM and associated with disease progression and prognosis ^41,42^ (Figure 4C).

This spatial bias implies a cis-regulatory mechanism, whereby mLINC00673 influences the expression of neighbouring genes. To test this, we investigated our gene regulatory network analysis and identified 60 lncRNAs with significant chromosomal bias in their inferred target genes (Supplementary Table 3). Notably, LINC00673 exhibited strong enrichment for targets located on human chromosome 17 (p < 10⁻⁵). We further mapped LINC00673-inferred target genes on human chromosome 17 and overlaid them with chromosome conformation capture (Hi-C) data from pancreatic cancer cell lines (PANC-1 cells). A significant overlap was observed between LINC00673 targets and a region of high chromatin interaction frequency on the long arm of chromosome 17 (Figure 4D), supporting a role for LINC00673 in organizing transcriptional hubs within this domain. Intriguingly, the distal portion of mouse chromosome 11 is syntenic with human chromosome 17 (Figure 4E), suggesting evolutionary conservation of LINC00673’s regulatory domain.

In summary, our data suggest that mLINC00673 regulates the expression of genes clustered within a syntenic region of mouse chromosome 11 and human chromosome 17 through long-range chromatin interactions. These genes, including cytokines critical for immune cell recruitment and tissue remodeling, underscore mLINC00673’s role as a nuclear scaffold for transcriptional regulation in pancreatic homeostasis and neoplastic transformation.

### LINC00673 facilitates RNA Polymerase II recruitment to proximal target genes

Recent studies have demonstrated that transcriptional activity from super-enhancers promotes the recruitment and stabilization of RNA Polymerase II (Pol II) at genes essential for cell lineage commitment (Figure 5A)^43,44^. To assess whether LINC00673 is required for Pol II localization at its putative target genes, we performed chromatin immunoprecipitation (ChIP) using an antibody specific for Pol II phosphorylated at serine 5 of its C-terminal domain (Pol II Ser5), a marker of transcriptional initiation. PANC-1 cells were transfected with either non-targeting control siRNA or siRNA targeting *LINC00673*, and efficient knockdown was confirmed by qRT-PCR (Figure 5B). Knockdown of *LINC00673* resulted in a significant reduction in Pol II Ser5 occupancy at the transcriptional start sites of *CCL8* and *RNF43*, which are located within the same genomic domain. These results indicate that LINC00673 is necessary for efficient recruitment of the transcriptional machinery to genes in its genomic vicinity.

**Figure 5:**
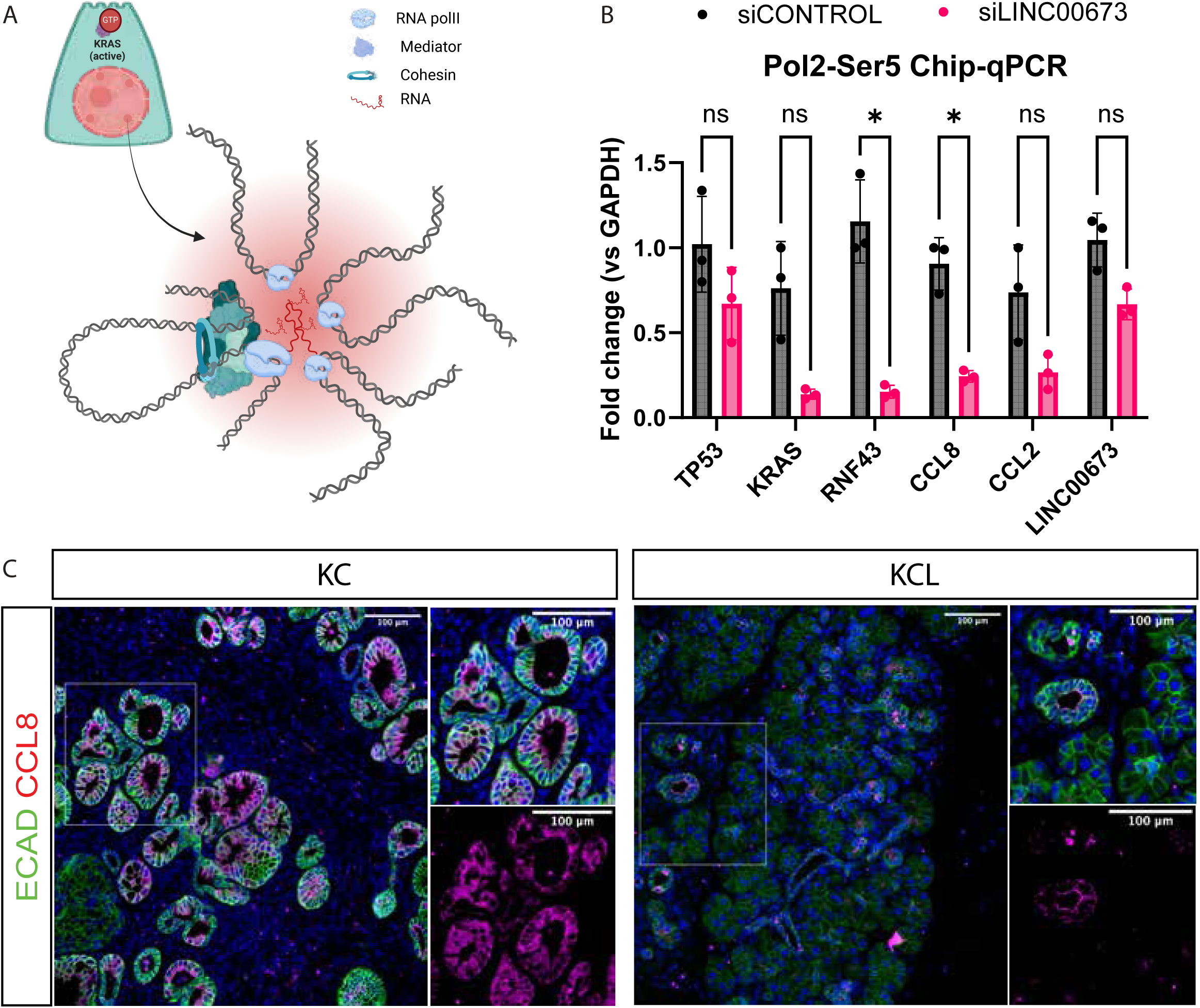
mLINC00673 controls the transcription machinery availability of its genomic vicinity. (A) Model: mLINC00673, transcribed from a super-enhancer, facilitates RNA Polymerase II (Pol II) recruitment and stabilization at nearby genes. (B) ChIP-qPCR for Pol II Ser5P in Panc1 cells following LINC00673 knockdown revealed reduced occupancy at target loci in the genomic vicinity. n = 3. Data are mean ± SD. *p < 0.05, **p < 0.01, ***p < 0.001. (C) Immunofluorescence staining of CCL8 (magenta) in preneoplastic lesions of PDAC (21 days post-injury with mutant Kras). Representative images from n = 3 mice.

Among these targets, CCL8 is a well-characterized chemoattractant for myeloid cells through CCR1 binding ^38^. Notably, CCL8 expression was significantly downregulated in PanIN lesions from KCL^HOM^ mice compared to KC controls 21 days after the induction of pancreatitis with CAE (Figure 5C), consistent with the reduced infiltration of immunosuppressive macrophages observed upon epithelial deletion of LINC00673 during early pancreatic tumorigenesis. Given that pharmacological inhibition of CCR1 has been shown to limit macrophage infiltration and promote tissue regeneration during PDAC initiation ^38^, this finding provides mechanistic insight into the inflammatory phenotype associated with LINC00673 loss.

Collectively, these data suggest that conditional deletion of mLINC00673 in epithelial cells impairs the recruitment of immunosuppressive macrophages, at least in part through the transcriptional downregulation of CCL8, thereby facilitating tissue repair. These findings highlight a central role for LINC00673 in coordinating immune-stromal interactions during early pancreatic lesion formation through regulation of a spatially confined pro-inflammatory gene network.

## Discussion

Significant advances in sequencing and computational methodologies have facilitated extensive integrative analyses of genomic data. Maps of open chromatin have unveiled clusters of regulatory regions within intergenic areas linked to developmental loci ^45,46^. Multiple studies, including our own, have demonstrated that these regulatory elements are extensively transcribed, yielding tissue-specific lncRNAs that are dynamically regulated during both development and disease ^47,48^. However, the specific functions of these transcribed elements remain elusive. In this study, we investigated the functional role of a transcribed regulatory element within the *Sox9* locus in the initiation of cancer. Our research uncovered that *LINC00673* is dynamically regulated during pancreas development and is induced in response to tissue injury and the expression of mutant Kras^G12D^ in the pancreas. Our data strongly indicate that *LINC00673* expression contributes to the creation of an oncogenic microenvironment that predisposes to cancer initiation by regulating the transcription of genes within its genomic vicinity.

Acinar dedifferentiation and ADM permit terminally differentiated, quiescent acinar cells to re-enter the cell cycle. Furthermore, acinar dedifferentiation leads to cytokine expression, tissue remodelling and the creation of a pro-inflammatory environment conducive to cancer development. These studies support the notion that cellular differentiation acts as a potent tumor suppressor, restraining both cell-autonomous and non-autonomous pro-tumorigenic processes in PDAC. Deletion of *mLINC00673* does not impede cellular proliferation and the upregulation of known ADM drivers such as *Sox9*, *Klf5* or *Gata6*, among others. Our data indicate that acinar dedifferentiation alone is insufficient for malignant transformation, suggesting that besides cellular dedifferentiation, additional cues are necessary for ADMs to progress into PanINs and PDAC. Cytokine activation and cell-cell communication are emerging as a pivotal feature of acinar dedifferentiation and tumor progression ^35^. Our data show that *mLINC00673* regulates the gene expression of a Ccl cytokine cluster located on chromosome 11. Among them *Ccl8* has been shown to be secreted by the epithelium and contributes to generating an immunosuppressive microenvironment and promote tumor progression ^38^. Our results dissociate cellular differentiation from cancer risk and underscore the importance of non-cell-autonomous processes and cell-cell communication in the transformation and progression of preinvasive PanIN lesions induced by KRAS. These findings hold significant implications regarding cancer risk and rejuvenating targeting strategies through the transient reprogramming of differentiated cells.

The genome is organized into insulated neighborhoods of highly interacting domains ^49–52^. This organizational structure is established during development, facilitating the emergence of cell-type-specific genome contacts and gene expression programs crucial for generating cellular diversity ^45,53–56^. LncRNAs often reside at the borders of these interacting domains within broad clusters of regulatory elements associated with development, suggesting their potential importance in genome organization and gene expression ^24,57–59^. In our study, we demonstrate that the deletion of exon 3 of *mLINC00673* leads to a disproportionate dysregulation of genes on the same chromosome. Recent studies have revealed that RNA molecules transcribed from enhancers play a role in regulating transcription at key developmental genes through RNA-protein interactions with Brd4, CTCF and the recruitment of other architectural proteins such as mediator and cohesins ^43,44^. Our findings propose that *mLINC00673* may regulate transcriptional processes through the activation of Pol2 to key developmental genes during injury-induced neoplastic transformation. Importantly, our gene targeting strategy preserves CTCF binding sites, implying that the recruitment may, at least in part, be mediated by the RNA itself. Future studies will focus on understanding how transcription from the *LINC00673* locus and the recruitment of regulatory elements coordinate genome organization and gene expression during inflammation-induced cancer initiation. Notably, the characteristics of other developmental enhancers resemble those of the LINC00673 locus, suggesting that LINC00673 may exemplify a general principle of genome regulation in both development and disease.

Our results demonstrate that cancer initiation depends on developmental regulatory elements that become activated during regeneration and can be exploited as novel therapies for this intractable disease. Moreover, our work implies that chromosomes are organized in functional units that operate beyond the definition of a gene and regulate, among other things, cell fate decisions with crucial biological impacts in development and cancer.

## Abbreviations

ADM: Acinar-to-ductal metaplasia
PDAC: pancreatic ductal adenocarcinoma
PanIN: pancreatic intraepithelial neoplasia
IPMN: intraductal papillary mucinous neoplasm
lncRNAs: long noncoding RNAs
Kras: Kirsten rat sarcoma proto-oncogene
ISH: *in situ* hybridization
hPSC: human pluripotent stem cells
CAE: caerulein
Hi-C: high-throughput chromosome conformation capture
scRNAseq: single cell RNAseq

## Author Contribution

Study concept and design: LA

Acquisition of data: MB, AK, HCM, EE, LA, PAS Drafting of the manuscript: MB, LA

Critical revision of the manuscript for important intellectual content: all authors Obtained funding: LA

Technical, or material support: HCM, EE, CVR, JH, PK, PV Study supervision: LA

## Acknowledgments

We would like to thank the Single Cell Genomics and Histology and Microscopy Core Facilities at the Biotech Research and Innovation Centre (BRIC) for their support and technical expertise.

## Disclosure

The authors declare that they have no conflicts of interest or financial relationships that could have influenced the research or the interpretation of the results.

## Materials and Methods

### Generation of a conditional knockout of mLINC00673 mouse line

The conditional allele of the exon3 of mLINC00673 was generated by homologous recombination as previously described ^60^ with few modifications. We flanked exon 3 of the mouse ortholog of *LINC00673*, *2610035d17Rik* (in this manuscript referred to as *mLINC00673*) with a loxP site (L83) and FNFL (FRT-Neo-FRT-LoxP) cassette which conferred G418 resistance. The targeting construct was generated on a bacterial artificial chromosome (BAC) with 8 kb- and 3 kb-long homology arms in the 5’end to L83 and 3’end to FLFL respectively (Supplementary Figure 4A). The vector also carried a diphtheria toxin alpha chain (DTA) negative selection marker to reduce random integration events. The vector was electroporated into PTL1 ESCs (129/B6 hybrid; Columbia University Transgenic Facilities) and after antibiotic selection we obtained two targeted ES cells identified by PCR screening. One was injected into C57BL/6 blastocysts to generate chimeric mice which were crossed with ACTB:FLPe mice (MGI:J:78660) to remove the FRT-flanked Neo cassette (Supplementary Figure 4B) and obtain a founder line of mice (mLINC00673) with the exon 3 flanked by loxP sites.

### Mouse models

LSL-KrasG12D (MGI:2429948), Pdx1Cre (MGI:2684321), R26RYFP (MGI:2449038), Ptf1aCreER (MGI:2387804) and mLINC00673fl mouse strains were used for breeding. Mice were maintained on a genetically mixed C57BL/6J-CD1 background. Animals were housed in accordance with best animal husbandry guideline recommendations of the European Union Directive (2010/63/EU). The Danish Animal Experiments Inspectorate reviewed and approved all animal experiments.

### In vivo mouse experiments

Recombination in *Ptf1a*^CreER^-bearing mice was induced by three intraperitoneal (i.p.) injections of 3 mg tamoxifen (T5648, Sigma-Aldrich) in corn oil (30 mg/ml) (C8267, Sigma Aldrich) administered every other day, over a period of five days. Injections were followed by a two-week washout period. To induce tissue injury in the pancreas, the cholecystokinin analogue caerulein (C9026; Sigma-Aldrich) was administered in PBS *via* i.p. injection every hour for seven hours (eight injections/day) on two consecutive days at a concentration of 125 mg/kg body weight as described elsewhere ^61^. Control mice were injected with an equivalent volume of PBS vehicle following the same protocol. Survival curve animals were tracked until the humane end point was met.

### Cell culture and maintenance

PANC-1 cells (ATCC; CRL-1469) were cultured in DMEM/GlutaMax media (Gibco, 31966-047) supplemented with 10% fetal bovine serum (FBS, Gibco, 17964671) and 1% penicillin-streptomycin (PenStrep, Gibco, 15-140-122). Cells were maintained at 37°C in a humidified atmosphere containing 4% CO₂.

### hESC differentiation

Hues8 cells were differentiated to pancreatic progenitors (PP) using an adapted version of ^28^. Cells were maintained in Nutristem (Biological Industries Israel Beit-Haemek Ltd., 05-100-1A) with daily media change and differentiation started when cells were confluent. Cells were cultured at 37 °C and 5% CO_2_. Cell passaging was performed with Accutase (Thermo Fisher Scientific, A1110501) in the embryonic stage or *via* TrypleE (Thermo Fisher Scientific, 12604013), Dispase (Sigma-Aldrich, D4693-1G) and mechanical dissociation in advanced stages of differentiation. When in suspension, cell media was supplemented with 10 μM ROCK inhibitor Y-27632 (VWR, 688000-100MG) that was maintained overnight when passaging.

Throughout Stage 1 and 4 cells were cultured in 2D with daily media change supplemented as stated in Supplementary Table 5. Then, cells were digested using TrypleE and grown in Geltrex (Fisher Scientific, A1413302) in PEm media (Multipotent pancreatic progenitors (MPPs) medium supplemented with 10 μM ROCK inhibitor). Differentiation protocol by ^28^.

### Patient-derived organoids

Fine-needle biopsy samples were grown in Geltrex (Fisher Scientific, A1413302) as previously described with minor changes ^62^. For passaging, organoids were mechanically dissociated with a 1 ml syringe, resuspended in 85% Geltrex and cultured in 7.5 µl domes.

### RNA extraction and cDNA synthesis of pancreatic tissue

RNA was extracted using RNeasy Mini Kit (Qiagen #74106) according to the manufacturer’s instructions for high RNase content tissues. Concentration and sample quality of the extracted RNA were measured using a spectrophotometer (DeNovix, DS-11). Retrieved RNA was treated with DNase to remove genomic DNA (Invitrogen #18068015). Samples were stored at −80°C. cDNA was synthesized using the GoScript Reverse Transcription System kit (Promega #A5001).

### Quantitative Reverse Transcription Polymerase Chain Reaction (qRT-PCR)

Expression levels were calculated using the comparative ddCT method of relative quantitation, with *Rpl5* (ribosomal protein L5) and *Rps29* (ribosomal protein S2) as housekeeping genes for mouse samples and *PPIA* (peptidylprolyl isomerase A) or *GAPDH* (Glyceraldehyde-3-Phosphate Dehydrogenase) for human samples (Supplementary Table 6).

### Immunohistochemistry and immunofluorescence

Pancreata were fixed in 4% paraformaldehyde (PFA) (9713-1000, VWR Chemicals) for 24 hours at room temperature and, after washing in PBS, were either immersed in 70% ethanol at room temperature (20824.365, VWR Chemicals) until paraffin embedding or 30% sucrose for cryoprotection in O.C.T. Tissues were cut into 4-10 μm sections and mounted on Superfrost Plus slides (10149870; Fisher Scientific). Antigen retrieval was performed with Tris-EDTA buffer at pH 9.0. Sections were washed in PBS-Tween 0.5% and blocked with 1% normal donkey serum before incubation with primary antibodies overnight at 4°C. Secondary antibodies were incubated with DAPI for 1 hour at room temperature. Sections were mounted with Vectashield® Mounting medium (H-1000; Vector Laboratories). Antibodies are listed in Supplementary Table 7.

H&E and alcian blue stainings were performed according to the manufacturer’s instructions (Abcam, ab245880 for H&E and Vector Laboratories, H-3501 for alcian blue) after which slides were mounted with Entellan (EMD Millipore, 1.07960.0500). Imaging was performed with a Hamamatzu NanoZoomer-XR Digital slide scanner C12000-01.

### Image acquisition

Immunofluorescence images were acquired *via* Olympus® ScanR screening. For quantification at least three animals were used per condition. For each animal, two sections of different depths within the pancreas were used with two distinct regions of each section being imaged. For each section, between 20 and 25 images were taken of each region, so 80-100 20x images were acquired per pancreas. Images with staining artifacts, lymph nodes or big vessels were deleted manually. Images in which a very low number of cells was detected were also discarded: an average of 500-3000 cells were detected within each of the selected images.

### Image quantification

For image quantification QuPath software (v.0.4.0) was used (https://qupath.github.io/). Area quantification using thresholder and cell classification using machine learning were performed according to standard protocols (https://qupath.github.io/).

For all quantifications two sections at different depths within the same pancreas were used. In the case of H&E and alcian blue quantifications whole-slide images were used. The brush tool was used to define an area of interest and exclude lymph nodes, spleen, big blood vessels, fat and staining artifacts.

For alcian blue area quantification color deconvolution was performed to correct the stain vectors using the built-in QuPath stain vector estimation in a representative area that included healthy acinar tissue, preneoplastic lesions and blank background. The resulting vectors were applied equally to all images. The average of channels was used to create a threshold classifier to select the epithelial tissue and remove clear interlobular infiltration. Next, a second thresholder was applied to the resulting annotation to detect the blue regions. Results were plotted as blue area (second threshold) as a percentage of the tissue area (first threshold).

Stroma quantification was performed using H&E staining and a supervised random tree-based classifier training to classify cells into stroma or epithelial cells. In brief, cells within the region of interest were detected using hematoxylin across all sections using the built-in cell detection tool. Then, regions of stromal or epithelial tissue were annotated and used to train a cell classifier. Additional annotations were added when needed. The percentage of stromal cells was then calculated per section and averaged between the two sections per pancreas. Stromal cells were represented as stromal cell percentage of total cells.

For immunofluorescence image quantification, a supervised random tree-based classifier training was used. First, cells were detected using DAPI with the built-in cell detection tool. Representative images of each tissue were used to classify positive and negative cells using the Point tool for each channel separately and used to train an object classifier based on all annotations. The classifiers of each channel were combined to generate a composite classifier to detect single-, double- and triple-positive cells that was applied to all the tiles. Successive round of training were performed as necessary. Results were plotted as indicated in the graphs.

### RNA in situ hybridization

Rnascope® Multiplex Fluorescent Reagent Kit v2 ACDBio (Newark, California, USA) was used following the manufacturer’s instructions. Paraffin sections were heated at 60°C for an hour, rehydrated, incubated with hydrogen peroxide and target retrieval was performed by boiling for 15 minutes. Protease IV 1:5 in RNase free water was applied for 15 minutes, and sections then incubated with RNAscope® Probe-Mm-2610035D17Rik (509101; ACDBio) or RNAscope™ Probe-Hs-LINC00673-O1 (548171; ACDBio) for 2 hours at. Subsequent immunofluorescence staining was perfomed as described above. Slides were mounted with Prolong® Gold Antifade Reagent (9071S; Cell Signaling Technology) and imaged *via* Olympus® ScanR screening or or with a Leica SP8 confocal microscope.

### RNA-sequencing and downstream analyses

Bulk RNA was sequenced with an Illumina NovaSeq 6000 Sequencing System with paired-end 150 bp read length and output of ≥ 20 million read pairs per sample (Novogene). Bioinformatic analysis was provided by Novogene.

### Knockdown and ChIP-qPCR

Cells were transfected using Lipofectamine™ RNAiMAX Transfection Reagent (Advanced Tech Group Inc, 13778150) according to the manufacturer’s protocol in conjunction with an siRNA targeting LINC00673 (siLINC00673, s500473). A non-targeting siRNA (siCONTROL, 4390844) served as negative control. Transfected cells were incubated for 48 hours before being harvested for downstream analyses. RNA was extracted using the RNeasy Mini Kit (Qiagen, 74106) following the manufacturer’s instructions. DNA was extracted as part of chromatin immunoprecipitation (ChIP) procedures detailed below. cDNA synthesis and quantitative PCR (qPCR) were performed as previously described. Chromatin immunoprecipitation (ChIP) was performed using the TruChip Chromatin Shearing Kit (Covaris, 520154) as previously described ^63^. In brief, cells were crosslinked with 16% formaldehyde, lysed, and chromatin was sheared to 100-500 bp fragment size with a Covaris E220 Ultrasonicator. DNA was immunoprecipitated with Phospho-RNA pol II CTD (Ser5) Monoclonal Antibody (Abcam, ab5131). Primer sequences are listed in Supplementary Table 6.

### Data availability

Raw RNA sequencing data will be uploaded into a repository upon manuscript publication. Accession numbers of publicly available data used in the manuscript are provided in Supplementary Table 8.

---

**Supplementary Figure 1:**
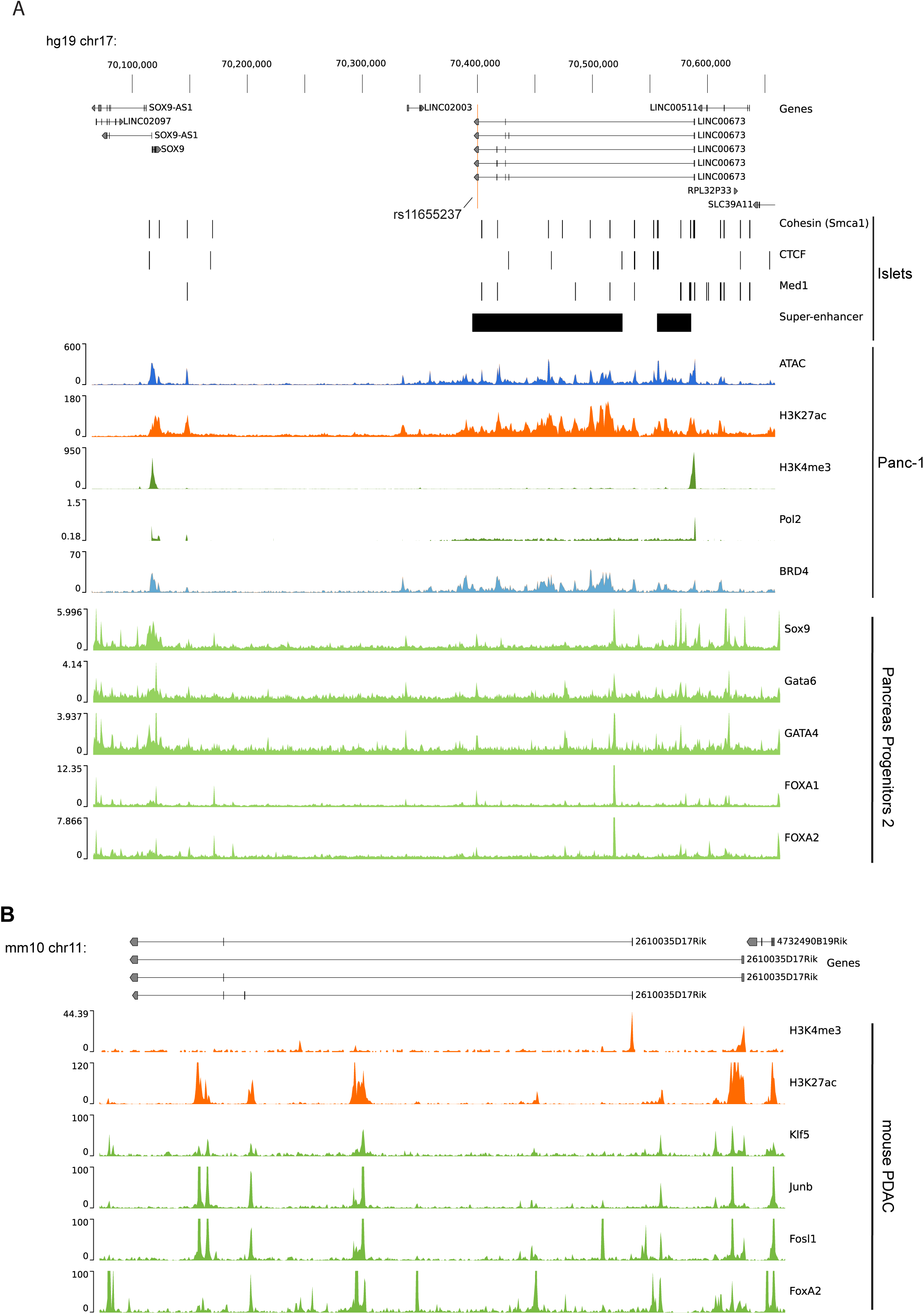
The LINC00673 locus has binding marks of TF of pancreas development, is transcribed from a super-enhancer and has marks of CTCF, Med1 and cohesins. A) Genome browser snapshot of the Sox9-LINC00673 locus on pancreatic lineages. LINC00673 harbors a SNP associated with PDAC survival (rs11655237) and is enriched in binding sites of cohesins (Smca1), CTCF and Med1 and is annotated as a super-enhancer in islets of Langerhans ^24^. Panc-1 ATAC-seq was obtained form ^64^, H3K27ac, H3K4me3 and Polymerase II ChIP-seq data were obtained from ENCODE ^65^, and BRD4 ChIP-seq from ^66^. B) Genome browser snapshot of the Sox9-mLINC00673 locus depicting H3K4me3, H3K27ac and transcription factor occupancy in tumor samples obtain from murine PDAC in the KPC model. H3K27ac, Klf5, Junb, Foxq1and FoxA2 ChiIP-seq of mouse pancreatic adenocarcinoma cancer cell line from^9^. H3K4me3 ChiIP-seq of mouse acinar cells from ^67^.

**Supplementary Figure 2.**
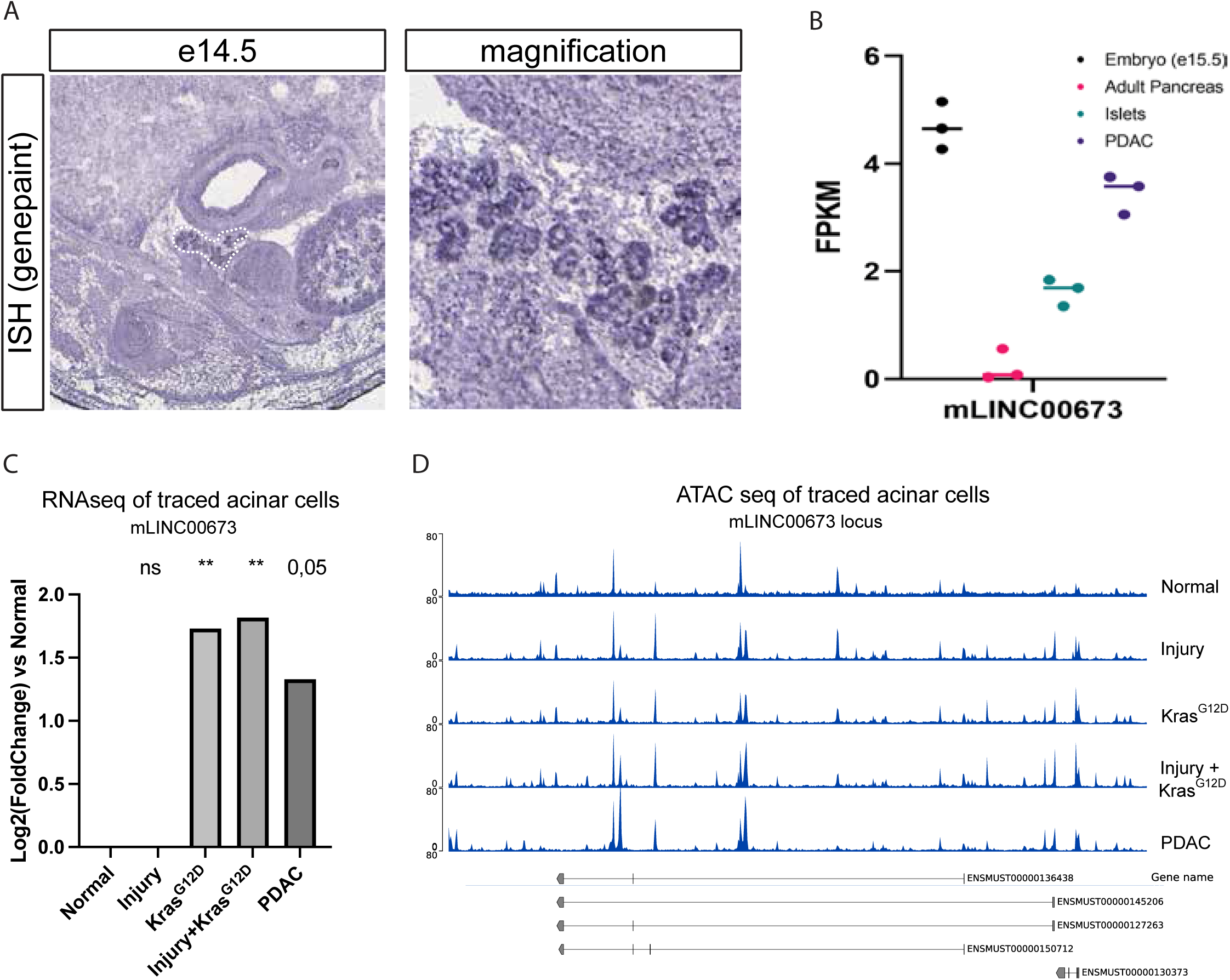
mLINC00673 expression is dynamically regulated in pancreas development and cancer. A) ISH of e14.5 mouse fetal pancreas (genepaint, https://gp3.mpg.de). B) We performed bulk RNA sequencing of mouse E15.5 embryo, adult pancreas, isolated islets and PDAC.N=3. C) We assessed chromatin accessibility (ATAC-seq) of the mLINC00673 locus of sorted lineage traced acinar cells. Data from ^27^. While the locus is accessible upon injury, expression is increased in KRAS-induced neoplastic transformation of the pancreatic epithelium.

**Supplementary Figure 3:**
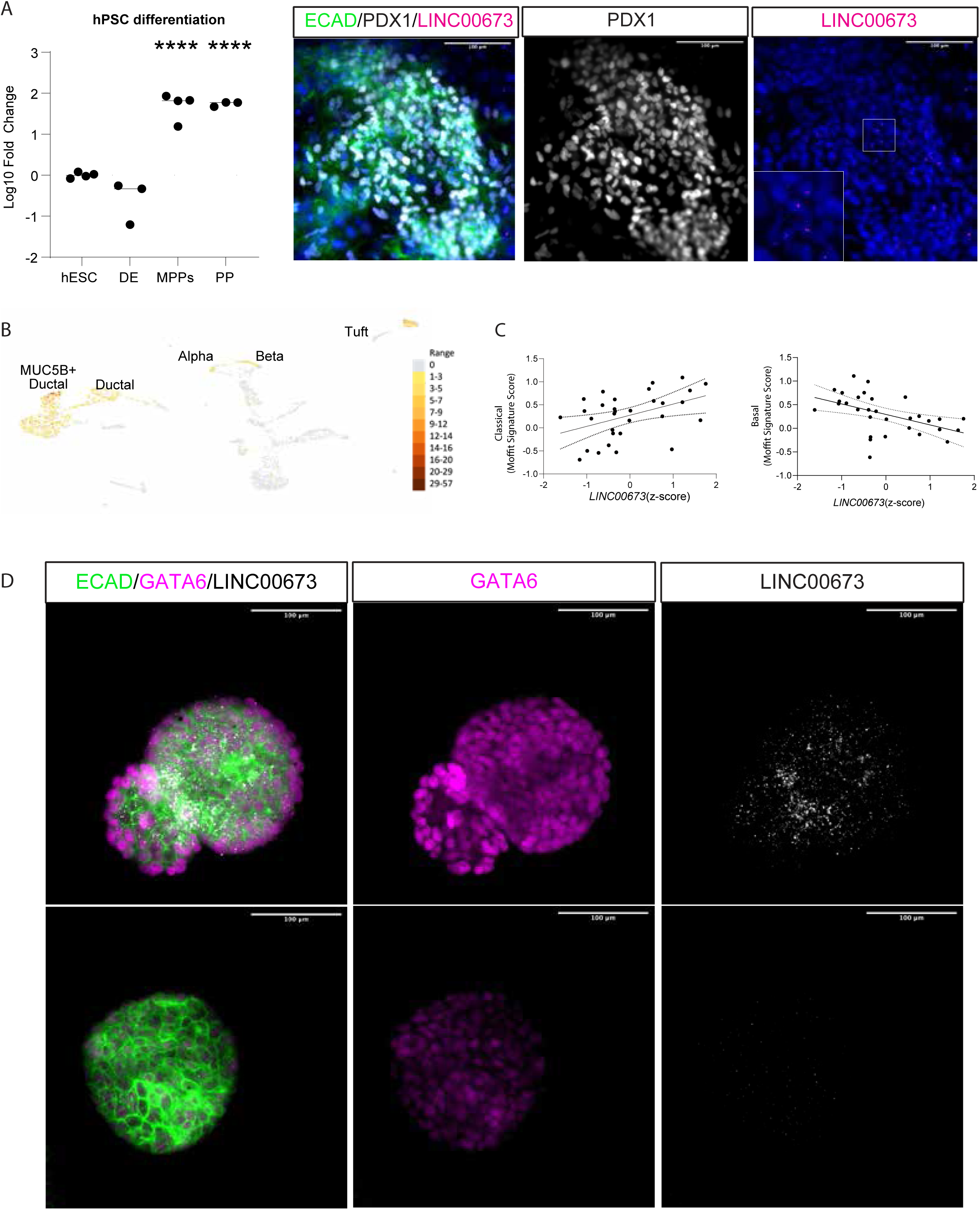
LINC00673 is expressed in pancreatic progenitors, ADM, and PDAC subtypes. A) Left: qPCR analysis of LINC00673 expression during in vitro differentiation of human pluripotent stem cells (hPSC) toward pancreatic progenitors. Expression peaked after the definitive endoderm stage (n = 3). Data are median; *p < 0.05, **p < 0.01, ****p < 0.001 by one-way ANOVA versus hESC. Right: RNAscope for LINC00673 in hPSC-derived pancreatic progenitors. B) Single-nucleus RNA-seq analysis of human chronic pancreatitis samples revealed LINC00673 expression in ADM cells (MUC5B+). Data from^30^. C) Left: LINC00673 expression in tumor-only (EPCAM+/CD45–) cells from resected PDAC samples correlated with classical and basal-like subtypes (n = 31). *p < 0.05, **p < 0.01, ***p < 0.005; R²_basal = 0.2448, R²_classical = 0.1903. Data from^31^. Right: RNAscope/IF in organoids from fine needle biopsies showed LINC00673 (white) expression correlates with high GATA6 (magenta) and E-cadherin (green) levels. Images acquired using identical laser settings (n = 3).

**Supplementary Figure 4:**
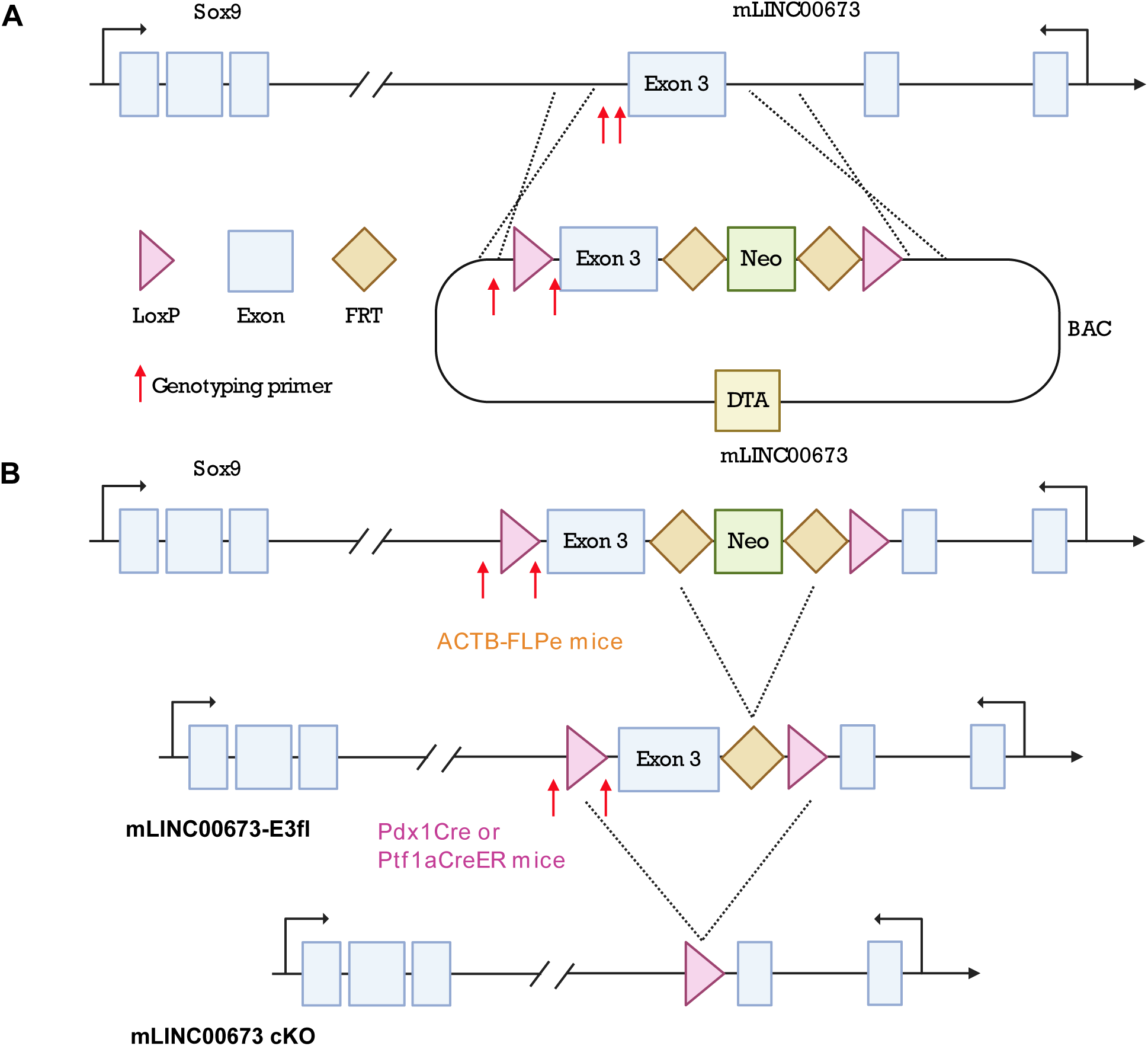
Generation of a conditional allele to study the function of mLINC00673 *in vivo*. (A) A targeting construct based on a BAC clone was used to flank exon 3 of mLINC00673 with loxP sites. (B) The resulting mice were crossed with ACTB-FLPe mice to excise the Neo cassette, generating mLINC00673^fl/fl^ animals, which were then crossed with Pdx1-Cre or Ptf1a-CreER mice for conditional deletion.

**Supplementary Figure 5.**
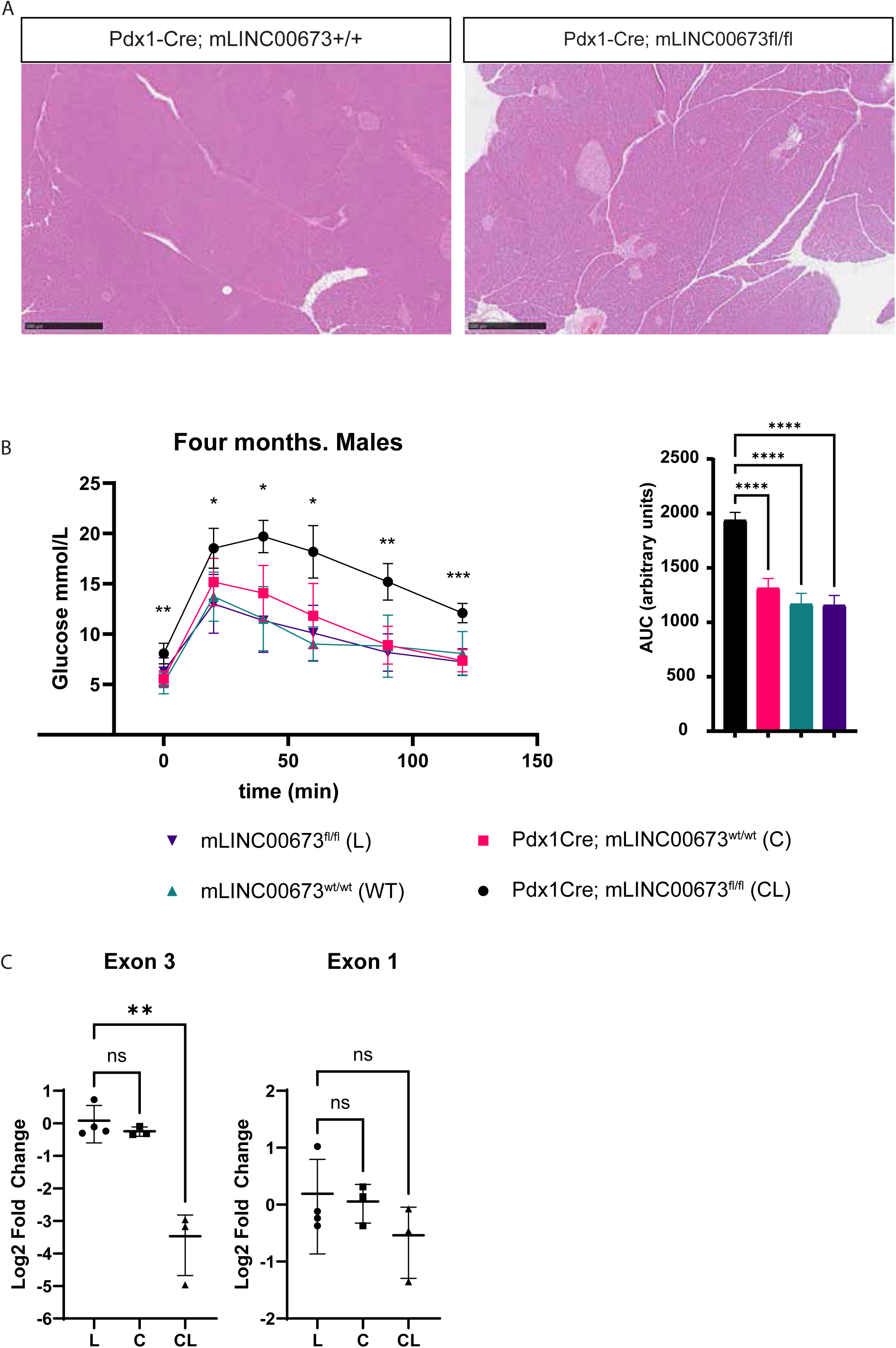
Characterization of mLINC00673 conditional knockout mice. (A) H&E staining of postnatal pancreas from control (C, CL) animals showed no morphological abnormalities. (B) Four-month-old mLINC00673 conditional knockout (cKO) males exhibited glucose intolerance in glucose tolerance tests; no significant changes were observed in females (n > 4). Right: area under the curve (AUC). Data are mean ± SD. *p < 0.05, **p < 0.01, ***p < 0.001 by one-way ANOVA with multiple comparisons. (C) qPCR analysis of E15.5 pancreata showed selective loss of exon 3 in Pdx1-Cre-driven knockouts, while exon 1 expression was preserved using exon-specific primers (n ≥ 3). *p < 0.05, **p < 0.01, ***p < 0.001 by one-way ANOVA.

**Supplementary Figure 6.**
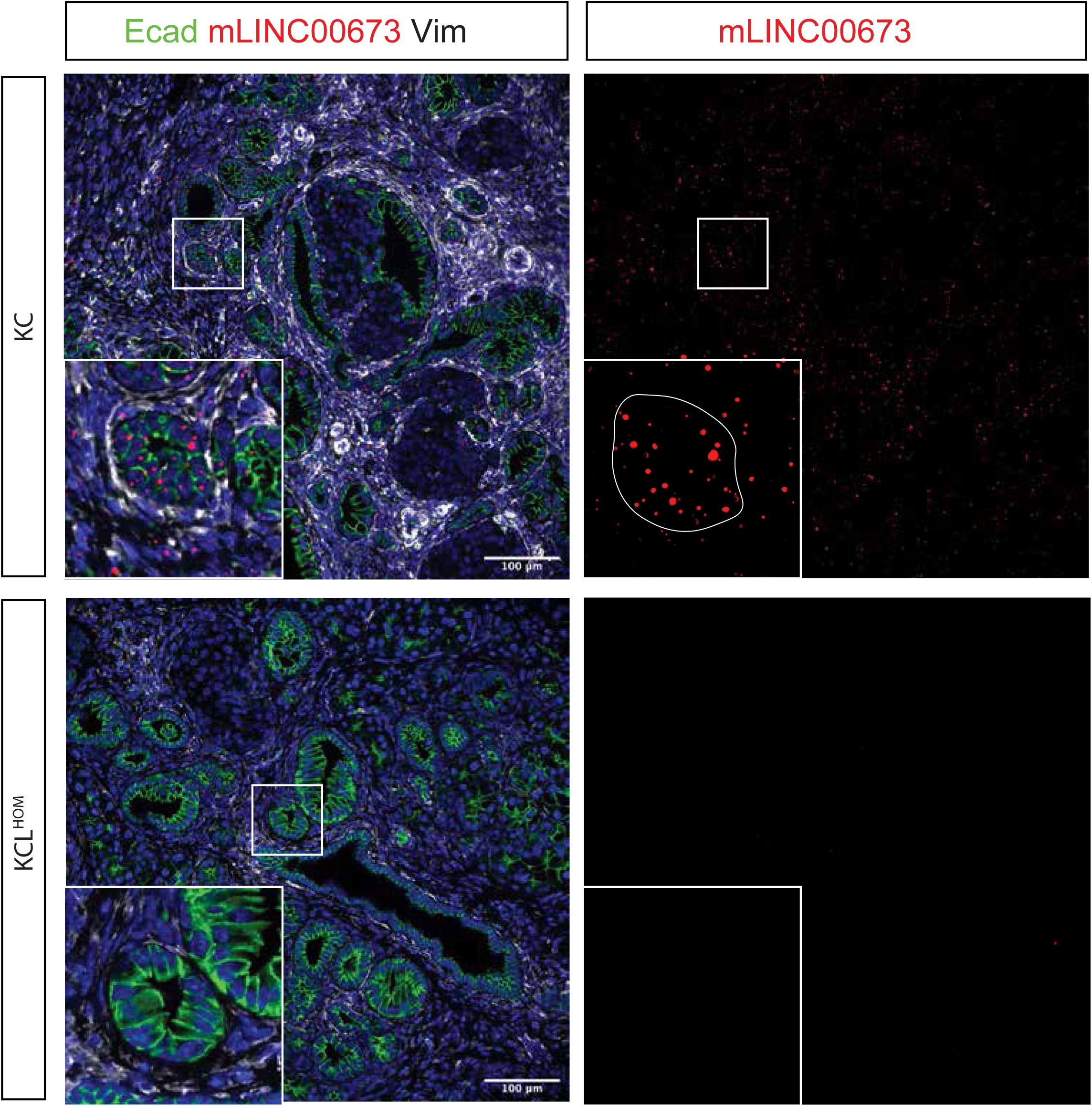
mLINC00673 expression is restricted to the epithelium in preneoplastic lesions. RNAscope for mLINC00673 (red) combined with immunofluorescence staining for E-cadherin (green) in pancreatic tissue 21 days after caerulein injections in KC and KCL^HOM^ mice (n = 3). mLINC00673 expression was confined to the epithelium in KC animals and absent in KCL^HOM^ mice. Images were acquired using identical laser intensity and exposure settings for both genotypes.

**Supplementary Figure 7.**
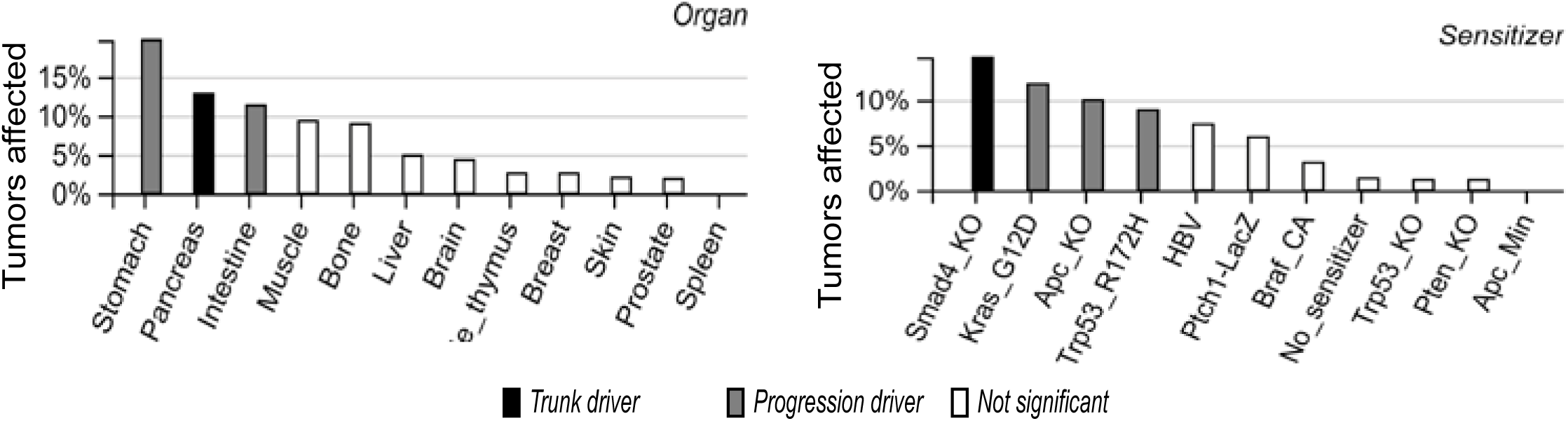
mLINC00673 acts as a trunk and progression driver in multiple tumor types. Analysis of the Sleeping Beauty cancer driver dataset identified mLINC00673 (2610035D17Rik) as both a trunk and progression driver in pancreatic, intestinal, and gastric tumors. The mLINC00673 locus was significantly altered in tumors driven by SMAD4 knockout, KRAS^G12D^, Trp53 loss, and activated WNT signaling.

**Supplementary Figure 8.**
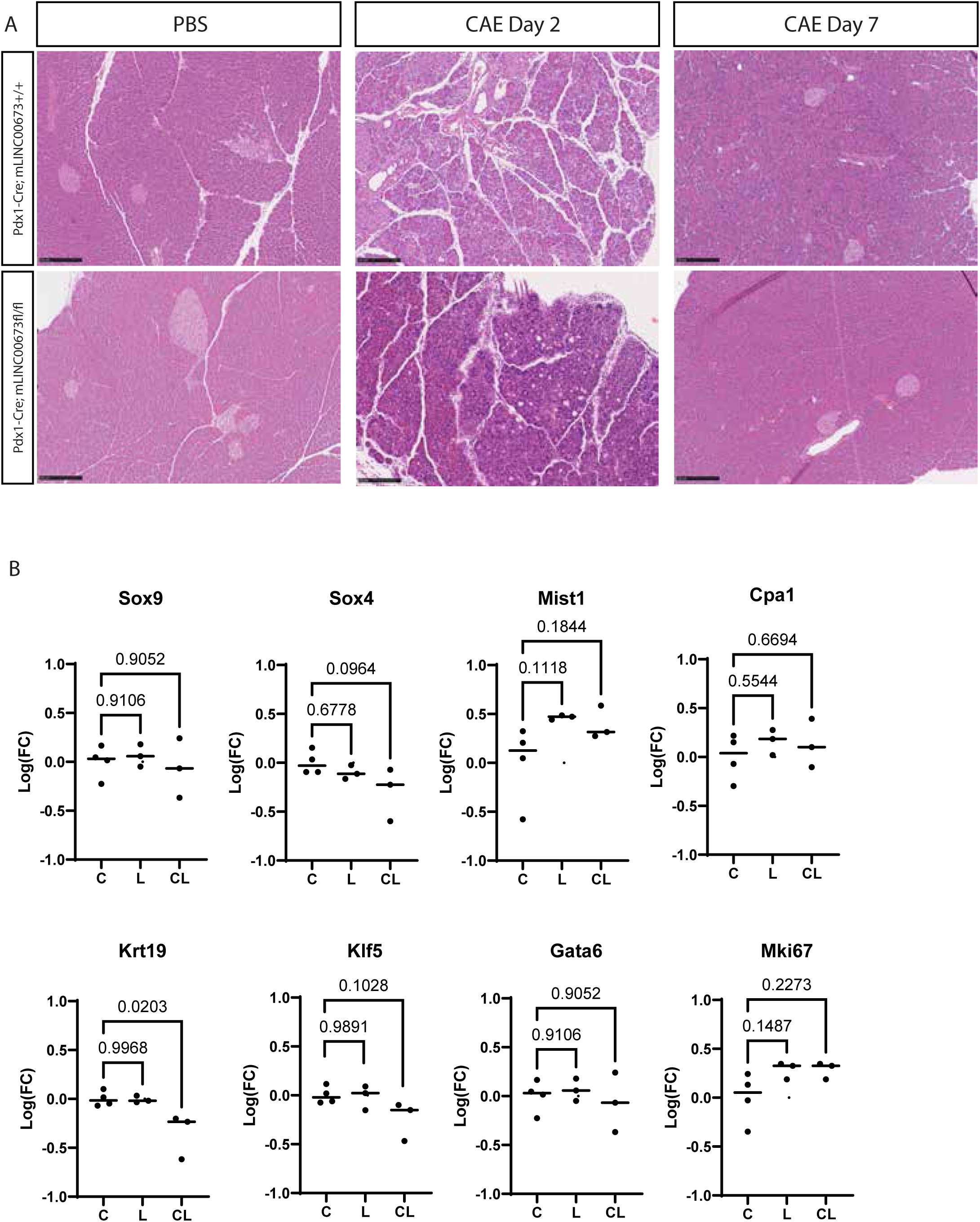
mLINC00673 is dispensable for pancreatic regeneration after acute injury. (A) Caerulein (CAE) was administered for two consecutive days to induce acute injury, and pancreata were harvested 2 days later. H&E staining showed no major differences in ADM or regeneration between WT and mLINC00673^KO^ mice (n = 3). (B) qPCR analysis of markers of acinar differentiation and plasticity revealed no significant impairment in mLINC00673^KO^ pancreata post-injury. Data are median; *p < 0.05 or as indicated in graphs by one-way ANOVA with multiple comparisons (n ≥ 3).

**Supplementary Figure 9:**
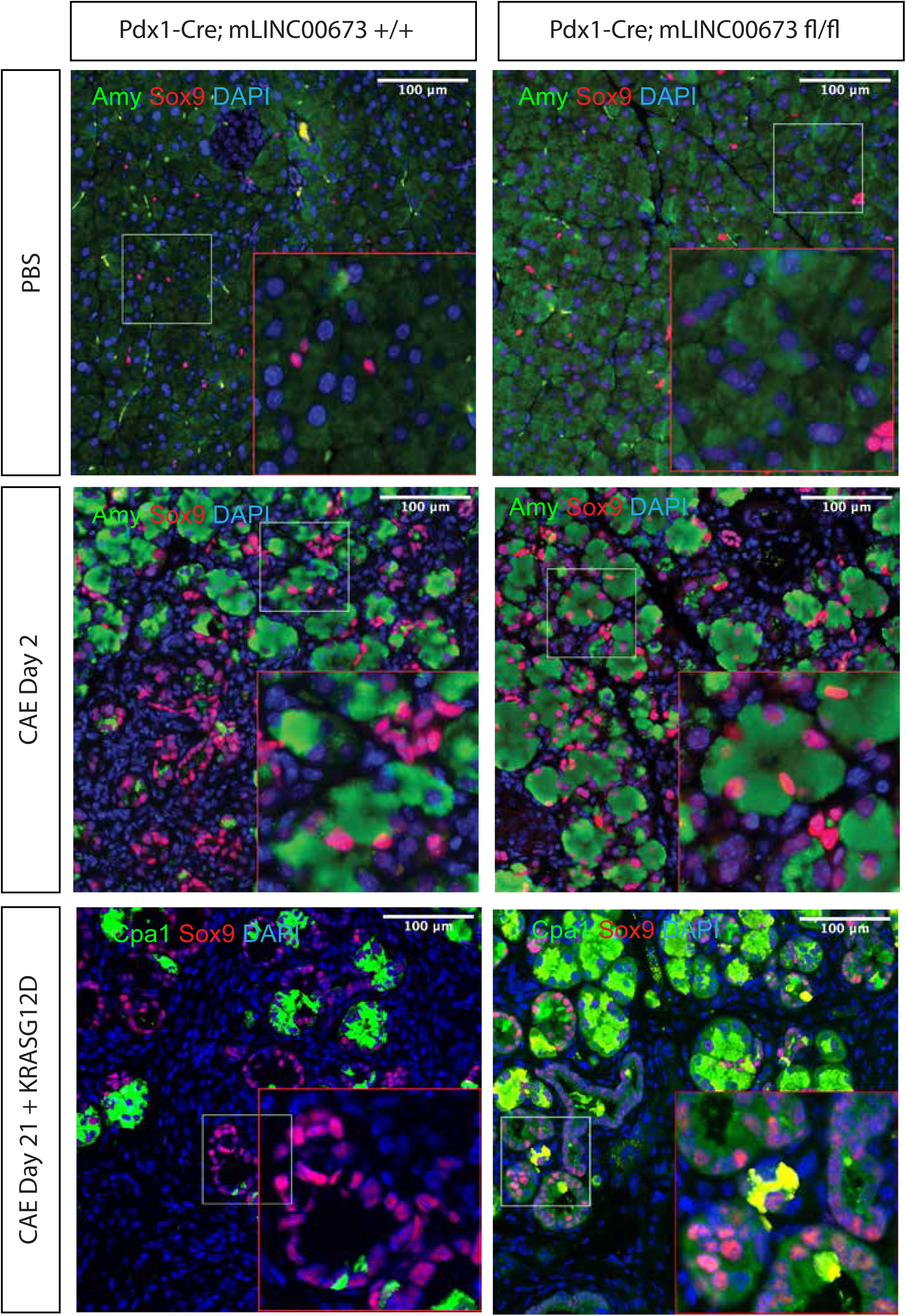
mLINC00673 does not regulate Sox9 expression. Immunofluorescence analysis of Sox9 (magenta) and either Amylase (Amy) or Cpa1 (green) in pancreata from control, caerulein-injured (2 days), and mutant Kras-expressing mice (Pdx1-Cre; mLINC00673^fl/fl^ and mLINC00673^cKO^). Homeostatic pancreas served as baseline control. Representative images from n = 3 animals.

**Supplementary Figure 10.**
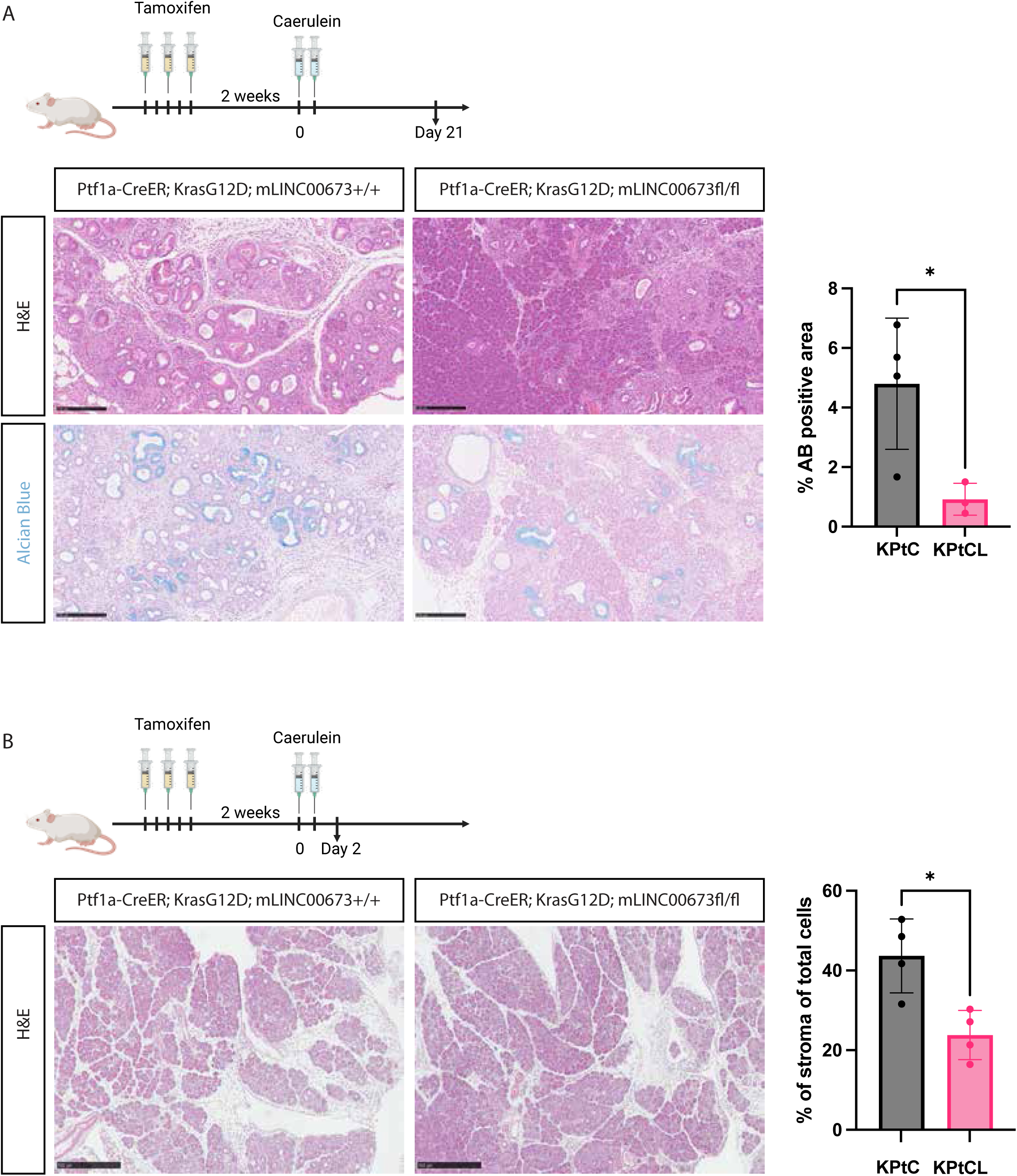
Acinar-specific deletion of mLINC00673 reduces PanIN formation and stromal infiltration. (A) To test the role of mLINC00673 in acinar-derived PanINs, we generated Ptf1a-CreER; Kras^G12D^; mLINC00673^+/+^ (KPtC) and Ptf1a-CreER; Kras^G12D^; mLINC00673^fl/fl^ (KPtCL) mice. Recombination was induced by tamoxifen (3 doses), followed by caerulein (CAE) for 2 days. Pancreata were collected 21 days later. H&E and alcian blue staining revealed reduced mucinous PanIN lesions (alcian blue+) in KPtCL mice. Right: quantification of alcian blue+ area as percentage of total tissue. *p < 0.05, unpaired t test. Data are mean ± SD. (B) Pancreata were also harvested 2 days post-CAE to assess responses to acute injury in the presence of oncogenic Kras. Deletion of mLINC00673 exon 3 in acinar cells did not impair ADM but reduced stromal infiltration. Right: quantification of stromal cells as percentage of total cells. *p < 0.05, unpaired t test. Data are mean ± SD.

## Notes

### Competing Interest Statement

The authors have declared no competing interest.

## References

1. Krah, N.M., De La O, J.-P., Swift, G.H., Hoang, C.Q., Willet, S.G., Chen Pan, F., Cash, G.M., Bronner, M.P., Wright, C.V., MacDonald, R.J., and Murtaugh, L.C. (2015). The acinar differentiation determinant PTF1A inhibits initiation of pancreatic ductal adenocarcinoma. eLife 4. 10.7554/elife.07125.

2. Cobo, I., Iglesias, M., Flandez, M., Verbeke, C., Del Pozo, N., Llorente, M., Lawlor, R., Luchini, C., Rusev, B., Scarpa, A., and Real, F.X. (2021). Epithelial Nr5a2 heterozygosity cooperates with mutant Kras in the development of pancreatic cystic lesions. J Pathol 253, 174–185. 10.1002/path.5570.

3. Mallen-St Clair, J., Soydaner-Azeloglu, R., Lee, K.E., Taylor, L., Livanos, A., Pylayeva-Gupta, Y., Miller, G., Margueron, R., Reinberg, D., and Bar-Sagi, D. (2012). EZH2 couples pancreatic regeneration to neoplastic progression. Genes & development 26, 439–444. 10.1101/gad.181800.111.

4. Miao, Z.F., Lewis, M.A., Cho, C.J., Adkins-Threats, M., Park, D., Brown, J.W., Sun, J.X., Burclaff, J.R., Kennedy, S., Lu, J., et al. (2020). A Dedicated Evolutionarily Conserved Molecular Network Licenses Differentiated Cells to Return to the Cell Cycle. Dev Cell 55, 178–194.e177. 10.1016/j.devcel.2020.07.005.

5. Ji, B., Tsou, L., Wang, H., Gaiser, S., Chang, D.Z., Daniluk, J., Bi, Y., Grote, T., Longnecker, D.S., and Logsdon, C.D. (2009). Ras Activity Levels Control the Development of Pancreatic Diseases. Gastroenterology 137, 1072–1082.e1076. 10.1053/j.gastro.2009.05.052.

6. Guerra, C., Collado, M., Navas, C., Alberto, Hernández-Porras, I., Cañamero, M., Rodriguez-Justo, M., Serrano, M., and Barbacid, M. (2011). Pancreatitis-Induced Inflammation Contributes to Pancreatic Cancer by Inhibiting Oncogene-Induced Senescence. Cancer Cell 19, 728–739. 10.1016/j.ccr.2011.05.011.

7. Wollny, D., Zhao, S., Everlien, I., Lun, X., Brunken, J., Brüne, D., Ziebell, F., Tabansky, I., Weichert, W., Marciniak-Czochra, A., and Martin-Villalba, A. (2016). Single-Cell Analysis Uncovers Clonal Acinar Cell Heterogeneity in the Adult Pancreas. Dev Cell 39, 289–301. 10.1016/j.devcel.2016.10.002.

8. Tata, A., Chow, R.D., and Tata, P.R. (2021). Epithelial cell plasticity: breaking boundaries and changing landscapes. EMBO Rep 22, e51921. 10.15252/embr.202051921.

9. Li, Y., He, Y., Peng, J., Su, Z., Li, Z., Zhang, B., Ma, J., Zhuo, M., Zou, D., Liu, X., et al. (2020). Mutant Kras co-opts a proto-oncogenic enhancer network in inflammation-induced metaplastic progenitor cells to initiate pancreatic cancer. Nature Cancer 2, 49–65. 10.1038/s43018-020-00134-z.

10. Andersson, R., and Sandelin, A. (2020). Determinants of enhancer and promoter activities of regulatory elements. Nat Rev Genet 21, 71–87. 10.1038/s41576-019-0173-8.

11. Hon, C.-C., Ramilowski, J.A., Harshbarger, J., Bertin, N., Rackham, O.J.L., Gough, J., Denisenko, E., Schmeier, S., Poulsen, T.M., Severin, J., et al. (2017). An atlas of human long non-coding RNAs with accurate 5′ ends. Nature 543, 199–204. 10.1038/nature21374.

12. Gil, N., and Ulitsky, I. (2018). Production of Spliced Long Noncoding RNAs Specifies Regions with Increased Enhancer Activity. Cell Syst 7, 537–547.e533. 10.1016/j.cels.2018.10.009.

13. Tan, J.Y., Smith, A.A.T., Ferreira da Silva, M., Matthey-Doret, C., Rueedi, R., Sönmez, R., Ding, D., Kutalik, Z., Bergmann, S., and Marques, A.C. (2017). cis-Acting Complex-Trait-Associated lincRNA Expression Correlates with Modulation of Chromosomal Architecture. Cell Reports 18, 2280–2288. 10.1016/j.celrep.2017.02.009.

14. Amaral, P.P., Leonardi, T., Han, N., Viré, E., Gascoigne, D.K., Arias-Carrasco, R., Büscher, M., Pandolfini, L., Zhang, A., Pluchino, S., et al. (2018). Genomic positional conservation identifies topological anchor point RNAs linked to developmental loci. Genome Biology 19, 32. 10.1186/s13059-018-1405-5.

15. Maurer, H.C., Curiel-Garcia, A., Holmstrom, S., Laise, P., Palermo, C.F., Sastra, S.A., Andren, A., Li, Z., LeLarge, T., Sagalovskiy, I., et al. (2023). Ras-dependent activation of BMAL2 regulates hypoxic metabolism in pancreatic cancer. bioRxiv, 2023.2003.2019.533333. 10.1101/2023.03.19.533333.

16. Lachmann, A., Giorgi, F.M., Lopez, G., and Califano, A. (2016). ARACNe-AP: gene network reverse engineering through adaptive partitioning inference of mutual information. Bioinformatics 32, 2233–2235. 10.1093/bioinformatics/btw216.

17. Kopp, J., von Figura, G., Mayes, E., Liu, F.-F., Claire, John, Fong, Akiyama, H., Christopher, Jensen, K., et al. (2012). Identification of Sox9-Dependent Acinar-to-Ductal Reprogramming as the Principal Mechanism for Initiation of Pancreatic Ductal Adenocarcinoma. Cancer Cell 22, 737–750. 10.1016/j.ccr.2012.10.025.

18. Milan, M., Balestrieri, C., Alfarano, G., Polletti, S., Prosperini, E., Spaggiari, P., Zerbi, A., Diaferia, G.R., and Natoli, G. (2019). FOXA2 controls the cis-regulatory networks of pancreatic cancer cells in a differentiation grade-specific manner. The EMBO journal 38, e102161. 10.15252/embj.2019102161.

19. de Andrés, M.P., Jackson, R.J., Felipe, I., Zagorac, S., Pilarsky, C., Schlitter, A.M., Martinez de Villareal, J., Jang, G.H., Costello, E., Gallinger, S., et al. (2023). GATA4 and GATA6 loss-of-expression is associated with extinction of the classical programme and poor outcome in pancreatic ductal adenocarcinoma. Gut 72, 535–548. 10.1136/gutjnl-2021-325803.

20. Kalisz, M., Bernardo, E., Beucher, A., Maestro, M.A., Del Pozo, N., Millán, I., Haeberle, L., Schlensog, M., Safi, S.A., Knoefel, W.T., et al. (2020). HNF1A recruits KDM6A to activate differentiated acinar cell programs that suppress pancreatic cancer. Embo j 39, e102808. 10.15252/embj.2019102808.

21. Klein, A.P., Wolpin, B.M., Risch, H.A., Stolzenberg-Solomon, R.Z., Mocci, E., Zhang, M., Canzian, F., Childs, E.J., Hoskins, J.W., Jermusyk, A., et al. (2018). Genome-wide meta-analysis identifies five new susceptibility loci for pancreatic cancer. Nature communications 9, 556. 10.1038/s41467-018-02942-5.

22. Necsulea, A., Soumillon, M., Warnefors, M., Liechti, A., Daish, T., Zeller, U., Baker, J.C., Grützner, F., and Kaessmann, H. (2014). The evolution of lncRNA repertoires and expression patterns in tetrapods. Nature 505, 635–640. 10.1038/nature12943.

23. Geusz, R.J., Wang, A., Chiou, J., Lancman, J.J., Wetton, N., Kefalopoulou, S., Wang, J., Qiu, Y., Yan, J., Aylward, A., et al. (2021). Pancreatic progenitor epigenome maps prioritize type 2 diabetes risk genes with roles in development. eLife 10, e59067. 10.7554/eLife.59067.

24. Miguel-Escalada, I., Bonàs-Guarch, S., Cebola, I., Ponsa-Cobas, J., Mendieta-Esteban, J., Atla, G., Javierre, B.M., Rolando, D.M.Y., Farabella, I., Morgan, C.C., et al. (2019). Human pancreatic islet three-dimensional chromatin architecture provides insights into the genetics of type 2 diabetes. Nature Genetics 51, 1137–1148. 10.1038/s41588-019-0457-0.

25. Srinivas, S., Watanabe, T., Lin, C.S., William, C.M., Tanabe, Y., Jessell, T.M., and Costantini, F. (2001). Cre reporter strains produced by targeted insertion of EYFP and ECFP into the ROSA26 locus. BMC developmental biology 1, 4. 10.1186/1471-213x-1-4.

26. Hingorani, S.R., Petricoin, E.F., Maitra, A., Rajapakse, V., King, C., Jacobetz, M.A., Ross, S., Conrads, T.P., Veenstra, T.D., Hitt, B.A., et al. (2003). Preinvasive and invasive ductal pancreatic cancer and its early detection in the mouse. Cancer cell 4, 437–450. S153561080300309X [pii].

27. Alonso-Curbelo, D., Ho, Y.-J., Burdziak, C., Maag, J.L.V., Morris, J.P., Chandwani, R., Chen, H.-A., Tsanov, K.M., Barriga, F.M., Luan, W., et al. (2021). A gene– environment-induced epigenetic program initiates tumorigenesis. Nature 590, 642–648. 10.1038/s41586-020-03147-x.

28. Ameri, J., Borup, R., Prawiro, C., Ramond, C., Schachter, K.A., Scharfmann, R., and Semb, H. (2017). Efficient Generation of Glucose-Responsive Beta Cells from Isolated GP2(+) Human Pancreatic Progenitors. Cell Rep 19, 36–49. 10.1016/j.celrep.2017.03.032.

29. Baldan, J., Houbracken, I., Rooman, I., and Bouwens, L. (2019). Adult human pancreatic acinar cells dedifferentiate into an embryonic progenitor-like state in 3D suspension culture. Scientific Reports 9, 4040. 10.1038/s41598-019-40481-1.

30. Tosti, L., Hang, Y., Debnath, O., Tiesmeyer, S., Trefzer, T., Steiger, K., Ten, F.W., Lukassen, S., Ballke, S., Kühl, A.A., et al. (2021). Single-Nucleus and In Situ RNA-Sequencing Reveal Cell Topographies in the Human Pancreas. Gastroenterology 160, 1330–1344.e1311. 10.1053/j.gastro.2020.11.010.

31. Espinet, E., Gu, Z., Imbusch, C.D., Giese, N.A., Buscher, M., Safavi, M., Weisenburger, S., Klein, C., Vogel, V., Falcone, M., et al. (2021). Aggressive PDACs Show Hypomethylation of Repetitive Elements and the Execution of an Intrinsic IFN Program Linked to a Ductal Cell of Origin. Cancer discovery 11, 638–659. 10.1158/2159-8290.CD-20-1202.

32. Childs, E.J., Mocci, E., Campa, D., Bracci, P.M., Gallinger, S., Goggins, M., Li, D., Neale, R.E., Olson, S.H., Scelo, G., et al. (2015). Common variation at 2p13.3, 3q29, 7p13 and 17q25.1 associated with susceptibility to pancreatic cancer. Nature Genetics 47, 911–916. 10.1038/ng.3341.

33. Newberg, J.Y., Mann, K.M., Mann, M.B., Jenkins, N.A., and Copeland, N.G. (2018). SBCDDB: Sleeping Beauty Cancer Driver Database for gene discovery in mouse models of human cancers. Nucleic Acids Res 46, D1011–D1017. 10.1093/nar/gkx956.

34. Pérez-Mancera, P.A., Rust, A.G., van der Weyden, L., Kristiansen, G., Li, A., Sarver, A.L., Silverstein, K.A., Grützmann, R., Aust, D., Rümmele, P., et al. (2012). The deubiquitinase USP9X suppresses pancreatic ductal adenocarcinoma. Nature 486, 266–270. 10.1038/nature11114.

35. Burdziak, C., Alonso-Curbelo, D., Walle, T., Reyes, J., Barriga, F.M., Haviv, D., Xie, Y., Zhao, Z., Zhao, C.J., Chen, H.A., et al. (2023). Epigenetic plasticity cooperates with cell-cell interactions to direct pancreatic tumorigenesis. Science 380, eadd5327. 10.1126/science.add5327.

36. Li, Y., He, Y., Peng, J., Su, Z., Li, Z., Zhang, B., Ma, J., Zhuo, M., Zou, D., Liu, X., et al. (2021). Mutant Kras co-opts a proto-oncogenic enhancer network in inflammation-induced metaplastic progenitor cells to initiate pancreatic cancer. Nature Cancer 2, 49–65. 10.1038/s43018-020-00134-z.

37. Arnes, L., Akerman, I., Balderes, D.A., Ferrer, J., and Sussel, L. (2016). βlinc1 encodes a long noncoding RNA that regulates islet β-cell formation and function. Genes & development 30, 502–507. 10.1101/gad.273821.115.

38. Zhang, Y., Lazarus, J., Steele, N.G., Yan, W., Lee, H.J., Nwosu, Z.C., Halbrook, C.J., Menjivar, R.E., Kemp, S.B., Sirihorachai, V.R., et al. (2020). Regulatory T-cell Depletion Alters the Tumor Microenvironment and Accelerates Pancreatic Carcinogenesis. Cancer discovery 10, 422–439. 10.1158/2159-8290.CD-19-0958.

39. Liou, G.-Y., Byrd, C.J., Storz, P., and Messex, J.K. (2024). Cytokine CCL9 Mediates Oncogenic KRAS-Induced Pancreatic Acinar-to-Ductal Metaplasia by Promoting Reactive Oxygen Species and Metalloproteinases. International Journal of Molecular Sciences 25, 4726.

40. Hosein, A.N., Dangol, G., Okumura, T., Roszik, J., Rajapakshe, K., Siemann, M., Zaid, M., Ghosh, B., Monberg, M., Guerrero, P.A., et al. (2022). Loss of Rnf43 Accelerates Kras-Mediated Neoplasia and Remodels the Tumor Immune Microenvironment in Pancreatic Adenocarcinoma. Gastroenterology 162, 1303–1318 e1318. 10.1053/j.gastro.2021.12.273.

41. Schussler, M.H., Skoudy, A., Ramaekers, F., and Real, F.X. (1992). Intermediate filaments as differentiation markers of normal pancreas and pancreas cancer. The American journal of pathology 140, 559–568.

42. Delgado-Coka, L.A., Roa-Peña, L., Babu, S., Horowitz, M., Petricoin, E.F., III, Matrisian, L.M., Blais, E.M., Marchenko, N., Allard, F.D., Akalin, A., et al. (2024). Keratin 17 is a prognostic and predictive biomarker in pancreatic ductal adenocarcinoma. American Journal of Clinical Pathology 162, 314–326. 10.1093/ajcp/aqae038.

43. Sabari, B.R., Dall’Agnese, A., Boija, A., Klein, I.A., Coffey, E.L., Shrinivas, K., Abraham, B.J., Hannett, N.M., Zamudio, A.V., Manteiga, J.C., et al. (2018). Coactivator condensation at super-enhancers links phase separation and gene control. Science 361. 10.1126/science.aar3958.

44. Cho, W.K., Spille, J.H., Hecht, M., Lee, C., Li, C., Grube, V., and Cisse, II (2018). Mediator and RNA polymerase II clusters associate in transcription-dependent condensates. Science 361, 412–415. 10.1126/science.aar4199.

45. Dowen, J.M., Fan, Z.P., Hnisz, D., Ren, G., Abraham, B.J., Zhang, L.N., Weintraub, A.S., Schujiers, J., Lee, T.I., Zhao, K., and Young, R.A. (2014). Control of cell identity genes occurs in insulated neighborhoods in mammalian chromosomes. Cell 159, 374–387. 10.1016/j.cell.2014.09.030.

46. Gaulton, K.J., Nammo, T., Pasquali, L., Simon, J.M., Giresi, P.G., Fogarty, M.P., Panhuis, T.M., Mieczkowski, P., Secchi, A., Bosco, D., et al. (2010). A map of open chromatin in human pancreatic islets. Nature Genetics 42, 255–259. 10.1038/ng.530.

47. Morán, I., Akerman, I., van de Bunt, M., Xie, R., Benazra, M., Nammo, T., Arnes, L., Nakić, N., García-Hurtado, J., Rodríguez-Seguí, S., et al. (2012). Human β cell transcriptome analysis uncovers lncRNAs that are tissue-specific, dynamically regulated, and abnormally expressed in type 2 diabetes. Cell metabolism 16, 435–448. 10.1016/j.cmet.2012.08.010.

48. Cabili, M.N., Trapnell, C., Goff, L., Koziol, M., Tazon-Vega, B., Regev, A., and Rinn, J.L. (2011). Integrative annotation of human large intergenic noncoding RNAs reveals global properties and specific subclasses. Genes & development 25, 1915–1927. 10.1101/gad.17446611.

49. Dixon, J.R., Selvaraj, S., Yue, F., Kim, A., Li, Y., Shen, Y., Hu, M., Liu, J.S., and Ren, B. (2012). Topological domains in mammalian genomes identified by analysis of chromatin interactions. Nature 485, 376–380. 10.1038/nature11082.

50. Nora, E.P., Lajoie, B.R., Schulz, E.G., Giorgetti, L., Okamoto, I., Servant, N., Piolot, T., van Berkum, N.L., Meisig, J., Sedat, J., et al. (2012). Spatial partitioning of the regulatory landscape of the X-inactivation centre. Nature 485, 381–385. 10.1038/nature11049.

51. Pope, B.D., Ryba, T., Dileep, V., Yue, F., Wu, W., Denas, O., Vera, D.L., Wang, Y., Hansen, R.S., Canfield, T.K., et al. (2014). Topologically associating domains are stable units of replication-timing regulation. Nature 515, 402–405. 10.1038/nature13986.

52. Symmons, O., Uslu, V.V., Tsujimura, T., Ruf, S., Nassari, S., Schwarzer, W., Ettwiller, L., and Spitz, F. (2014). Functional and topological characteristics of mammalian regulatory domains. Genome research 24, 390–400. 10.1101/gr.163519.113.

53. Sima, J., Chakraborty, A., Dileep, V., Michalski, M., Klein, K.N., Holcomb, N.P., Turner, J.L., Paulsen, M.T., Rivera-Mulia, J.C., Trevilla-Garcia, C., et al. (2019). Identifying cis Elements for Spatiotemporal Control of Mammalian DNA Replication. Cell 176, 816–830.e818. 10.1016/j.cell.2018.11.036.

54. Stewart-Morgan, K.R., Reverón-Gómez, N., and Groth, A. (2019). Transcription Restart Establishes Chromatin Accessibility after DNA Replication. Mol Cell 75, 284–297.e286. 10.1016/j.molcel.2019.04.033.

55. Stadhouders, R., Vidal, E., Serra, F., Di Stefano, B., Le Dily, F., Quilez, J., Gomez, A., Collombet, S., Berenguer, C., Cuartero, Y., et al. (2018). Transcription factors orchestrate dynamic interplay between genome topology and gene regulation during cell reprogramming. Nature genetics 50, 238–249. 10.1038/s41588-017-0030-7.

56. Beagrie, R.A., Scialdone, A., Schueler, M., Kraemer, D.C.A., Chotalia, M., Xie, S.Q., Barbieri, M., de Santiago, I., Lavitas, L.-M., Branco, M.R., et al. (2017). Complex multi-enhancer contacts captured by genome architecture mapping. Nature 543, 519–524. 10.1038/nature21411.

57. Symmons, O., Pan, L., Remeseiro, S., Aktas, T., Klein, F., Huber, W., and Spitz, F. (2016). The Shh Topological Domain Facilitates the Action of Remote Enhancers by Reducing the Effects of Genomic Distances. Developmental cell 39, 529–543. 10.1016/j.devcel.2016.10.015.

58. Lupianez, D.G., Kraft, K., Heinrich, V., Krawitz, P., Brancati, F., Klopocki, E., Horn, D., Kayserili, H., Opitz, J.M., Laxova, R., et al. (2015). Disruptions of topological chromatin domains cause pathogenic rewiring of gene-enhancer interactions. Cell 161, 1012–1025. 10.1016/j.cell.2015.04.004.

59. Hnisz, D., Abraham, B.J., Lee, T.I., Lau, A., Saint-Andre, V., Sigova, A.A., Hoke, H.A., and Young, R.A. (2013). Super-enhancers in the control of cell identity and disease. Cell 155, 934–947. 10.1016/j.cell.2013.09.053.

60. Mastracci, T.L., Lin, C.-S., and Sussel, L. (2013). Generation of mice encoding a conditional allele of Nkx2.2. Transgenic Research 22, 965–972. 10.1007/s11248-013-9700-0.

61. Kong, B., Bruns, P., Behler, N.A., Chang, L., Schlitter, A.M., Cao, J., Gewies, A., Ruland, J., Fritzsche, S., Valkovskaya, N., et al. (2018). Dynamic landscape of pancreatic carcinogenesis reveals early molecular networks of malignancy. Gut 67, 146–156. 10.1136/gutjnl-2015-310913.

62. Driehuis, E., Gracanin, A., Vries, R.G.J., Clevers, H., and Boj, S.F. (2020). Establishment of Pancreatic Organoids from Normal Tissue and Tumors. STAR Protocols 1, 100192. 10.1016/j.xpro.2020.100192.

63. Orlando, David A., Chen, Mei W., Brown, Victoria E., Solanki, S., Choi, Yoon J., Olson, Eric R., Fritz, Christian C., Bradner, James E., and Guenther, Matthew G. (2014). Quantitative ChIP-Seq Normalization Reveals Global Modulation of the Epigenome. Cell Reports 9, 1163–1170. 10.1016/j.celrep.2014.10.018.

64. Ren, B., Yang, J., Wang, C., Yang, G., Wang, H., Chen, Y., Xu, R., Fan, X., You, L., Zhang, T., and Zhao, Y. (2021). High-resolution Hi-C maps highlight multiscale 3D epigenome reprogramming during pancreatic cancer metastasis. Journal of Hematology & Oncology 14, 120. 10.1186/s13045-021-01131-0.

65. Dunham, I., Kundaje, A., Aldred, S.F., Collins, P.J., Davis, C.A., Doyle, F., Epstein, C.B., Frietze, S., Harrow, J., Kaul, R., et al. (2012). An integrated encyclopedia of DNA elements in the human genome. Nature 489, 57–74. 10.1038/nature11247.

66. Hamdan, F.H., and Johnsen, S.A. (2018). DeltaNp63-dependent super enhancers define molecular identity in pancreatic cancer by an interconnected transcription factor network. Proc Natl Acad Sci U S A 115, E12343–e12352. 10.1073/pnas.1812915116.

67. Mehmood, R., Varga, G., Mohanty, S.Q., Laing, S.W., Lu, Y., Johnson, C.L., Kharitonenkov, A., and Pin, C.L. (2014). Epigenetic reprogramming in Mist1(-/-) mice predicts the molecular response to cerulein-induced pancreatitis. PloS one 9, e84182. 10.1371/journal.pone.0084182.

